# β-adrenergic control of sarcolemmal Ca_V_1.2 abundance by small GTPase Rab proteins

**DOI:** 10.1101/2020.09.28.317362

**Authors:** Silvia G. del Villar, Taylor L. Voelker, Heather C. Spooner, Eamonn J. Dickson, Rose E. Dixon

**Affiliations:** Dept. of Physiology and Membrane Biology, University of California, Davis, CA

**Keywords:** L-type calcium channel, heart, β-adrenergic receptor, trafficking, clustering

## Abstract

The number and activity of Ca_v_1.2 channels in the cardiomyocyte sarcolemma tunes the magnitude of Ca^2+^-induced Ca^2+^ release and myocardial contraction. β-adrenergic receptor (*βAR*) activation stimulates sarcolemmal insertion of Ca_V_1.2 channels. This supplements the pre-existing sarcolemmal Ca_V_1.2 population, forming large ‘super-clusters’ wherein neighboring channels undergo enhanced cooperative-gating behavior, amplifying Ca^2+^ influx and myocardial contractility. Here, we determine this stimulated insertion is fueled by an internal reserve of early- and recycling endosome-localized, pre-synthesized Ca_V_1.2 channels. *βAR*-activation decreased Ca_V_1.2/endosome colocalization in ventricular myocytes, as it triggered ‘emptying’ of endosomal Ca_V_1.2 cargo into the sarcolemma. We examined the rapid dynamics of this stimulated insertion process with live-myocyte imaging of channel trafficking, and discovered that Ca_V_1.2 are often inserted into the sarcolemma as pre-formed, multi-channel clusters. Likewise, entire clusters were removed from the sarcolemma during endocytosis, while in other cases, a more incremental process suggested removal of individual channels. The amplitude of the stimulated insertion response was doubled by co-expression of constitutively-active Rab4a, halved by co-expression of dominant-negative Rab11a, and abolished by co-expression of dominant-negative mutant Rab4a. In ventricular myocytes, *βAR*-stimulated recycling of Ca_V_1.2 was diminished by both nocodazole and latrunculin-A, suggesting an essential role of the cytoskeleton in this process. Functionally, cytoskeletal disruptors prevented *βAR-activated* Ca^2+^ current augmentation. Moreover, *βAR*-regulation of Ca_V_1.2 was abolished when recycling was halted by co-application of nocodazole and latrunculin-A. These findings reveal that *βAR*-stimulation triggers an on-demand boost in sarcolemmal Ca_V_1.2 abundance via targeted, Rab4a and Rab11a-dependent insertion of Ca_V_1.2 channels is essential for *βAR*-regulation of cardiac Ca_V_1.2.

**Significance Statement:** The L-type voltage-gated Ca^2+^ channel Ca_V_1.2 is essential for excitation-contraction coupling in the heart. During the fight-or-flight response, Ca_V_1.2 channel activity is augmented as a result of PKA-mediated phosphorylation, downstream of β-adrenergic receptor (*βAR*) activation. We discovered that enhanced sarcolemmal abundance of Ca_V_1.2 channels, driven by stimulated insertion/recycling of specific Ca_V_1.2 containing endosomes, is essential for *βAR*-mediated regulation of these channels in the heart. These data reveal a new conceptual framework of this critical and robust pathway for on-demand tuning of cardiac EC-coupling during fight-or-flight.

## Introduction

Ca^2+^ influx through L-type Ca^2+^ channels (Ca_V_1.2) is indispensable for cardiac excitation-contraction coupling (EC-coupling) (1). These multimeric proteins consist of a pore-forming and voltage-sensing α_1c_ subunit, and auxiliary β and α_2_δ-subunits. In ventricular myocytes, Ca_V_1.2 mainly localize to the t-tubule sarcolemma and open briefly, allowing a small amount of Ca^2+^ influx, in response to the wave of depolarization that travels through the conduction system of the heart from its point of origin, usually in the SA-node. This initial influx is amplified manifold though Ca^2+^-induced Ca^2+^ release (CICR) from juxtaposed type 2 ryanodine receptors (RyR2) on the junctional sarcoplasmic reticulum (jSR), Which are localized a short ~12 nm across the dyadic cleft. The synchronous opening of thousands of RyR2 generates a transient, global elevation in intracellular calcium concentration ([Ca^2+^]_i_), resulting in contraction. Reducing Ca_V_1.2 channel current (*I*_Ca_) results in less CICR, smaller [Ca^2+^]_i_ transients, and less forceful contractions. Conversely, larger amplitude *I*_Ca_ elicits greater Ca^2+^ release from the SR, producing more forceful contractions. The level of Ca^2+^ influx through Ca_V_1.2 channels therefore tunes EC-coupling.

*I*_Ca_ is a product of the number of channels in the sarcolemma and their open probability (*P*_o_). Consequently, there are two possible, non-mutually exclusive strategies that may be adopted to alter *I*_Ca_ and consequently the magnitude of EC-coupling: i) adjust Ca_V_1.2 channel activity (*P*_o_) and/or, ii) modify sarcolemmal Ca_V_1.2 channel expression (*N*). The first strategy of increasing channel *P*_o_ has long been associated with β-adrenergic receptor (*βAR*)-mediated signaling in the heart (2–4). During acute physical or emotional stress, norepinephrine spills from sympathetic varicosities onto cardiomyocytes, activating *βAR*s. The ensuing G_αs_/adenylyl cyclase/cAMP/PKA signaling cascade culminates in PKA phosphorylation of several effector proteins, including Ca_V_1.2 (or an element of their interactome (5, 6)), enhancing their activity to generate this positive inotropic response.

As to the second strategy to increase *I*_Ca_, there remains a paucity of information regarding the mechanisms regulating Ca_V_1.2 channel abundance in the cardiomyocyte sarcolemma. Classical secretory transport literature suggests that Ca_V_1.2 channels are trafficked from the endoplasmic reticulum (ER) to the trans-Golgi-network (TGN) and onward to their dyadic position in the sarcolemma. Underscoring the importance of faithful Ca_V_1.2 channel trafficking, altered Ca_V_1.2 channel density has been reported in both failing (7, 8) and aging (9) ventricular myocytes, and impaired anterograde trafficking of Ca_V_1.2 channels to the t-tubules of human ventricular myocytes has been linked to dilated cardiomyopathy (10). Yet, despite the importance of tight homeostatic control of Ca_V_1.2 channel trafficking to prevent Ca^2+^ dysregulation, the molecular steps defining Ca_V_1.2 channel sorting and insertion remain poorly understood. Therefore, elucidation of the trafficking pathways that regulate Ca_V_1.2 channel abundance is critical for our understanding of the pathophysiology of heart failure and myocardial aging, and could potentially reveal new therapeutic or rejuvenation targets. Along that vein, in the treatment of cystic fibrosis, multiple drugs are in various stages of use or development to improve trafficking to, or to amplify, or stabilize, CFTR channels at the apical membrane of airway epithelial cells (11).

There have been no measurements of Ca_V_1.2 channel lifetimes in cardiomyocytes, but pulse-chase experiments in immortalized cell lines support a lifetime of plasma membrane (PM)-localized Ca_V_1.2 channels of ~3 h (12) while total cellular Ca_V_1.2 lifetime is >20 h (13, 14). This disparity suggests membrane-Ca_V_1.2 turns over much more dynamically than the total cellular channel content and implies ongoing local control by endosomal trafficking. Disturbance of the equilibrium between channel insertion/recycling and internalization would be predicted to lead to alterations in sarcolemmal Ca_V_1.2 channel abundance. Trafficking of vesicular cargo through the endosomal pathway is regulated by Rab-GTPases, a >60-member family within the larger Ras superfamily of small GTPases (15–18). Rab5 is involved in endocytosis and control of vesicular cargo influx into early endosomes (EEs; also called sorting endosomes), while Rab4 controls efflux of cargo out of EEs and fast recycling (t_1/2_ ~1-2 min) back to the PM (19). Rab11, expressed on recycling endosomes (RE; also called the endocytic recycling compartment or ERC), regulates slow recycling (t_1/2_ ~12 min) of cargo from this compartment back to the PM (19). In cortical neurons and pancreatic β-cells, activity-dependent Ca_V_1.2 channel internalization has been postulated to play important roles in Ca^2+^ homeostasis, with implications for homeostatic synaptic plasticity and insulin production respectively (20). In mouse neonatal cardiomyocytes, Rab11b has been reported to limit expression PM Ca_V_1.2 (21), while recent studies performed in HEK and HL-1 cells reported that endocytic recycling of cardiac Ca_V_1.2 channels, regulates their surface abundance (13, 22). Despite this crucial information from other cell-types, there has been a lack of rigorous investigations, at the molecular level, into how Ca_V_1.2 channel recycling is regulated in cardiac myocytes.

Here, we identify a dynamic, sub-sarcolemmal pool of Ca_V_1.2-cargo-carrying endosomes that are rapidly mobilized to the ventricular myocyte sarcolemma along targeted Rab4a and Rab11a GTPase-regulated recycling pathways in response to βAR-stimulation. Using electrophysiology, cell biology approaches, TIRF and super-resolution microscopy, we report that enhanced sarcolemmal Ca_V_1.2 abundance via targeted, ISO-stimulated, recycling of Ca_V_1.2 channels is essential for *β*AR-regulation of cardiac Ca_V_1.2.

## Results

### Internal pools of pre-synthesized Ca_V_1.2 channels reside on endosomes

Recently, we reported that sarcolemmal Ca_V_1.2 channel expression increases in the heart during *βAR* signaling (23). Activation of *βAR*s with isoproterenol (ISO) produced a rapid, PKA-dependent augmentation of Ca_V_1.2 channel abundance along ventricular myocyte t-tubules. We hypothesized that an endosomal pool of Ca_V_1.2 fuels the rapid, ISO-stimulated insertion of channels into the sarcolemma of ventricular myocytes. To test this, we performed an examination of the distribution of Ca_V_1.2 channels on EEs, REs and late endosomes (LEs) in adult mouse ventricular myocytes (AMVMs) using two- or three-color Airyscan super-resolution microscopy. Immuno-stained Ca_V_1.2 channel cargo was observed on 15.1 ± 0.4 % of early endosome antigen-1 (EEA1) positive pixels in male ventricular myocytes and on a similar 16.1 ± 0.7 % in female myocytes (Figure 1A and B; *P* = 0.24; see also Figure S1). Pre-fixation stimulation with 100 nM ISO led to a significant, 15-20 % decrease in colocalization between Ca_V_1.2 and EEA1 (male *P* = 0.001, female *P* = 0.01). We further narrowed our analysis to Rab4 positive EEs by performing triple-label experiments, co-staining for Ca_V_1.2, EEA1, and Rab4 (Figure S2). A similar trend was observed there such that 15.6 ± 1.3 % of EEA1- and Rab4 co-expressing pixels were colocalized with Ca_V_1.2, falling to 10.8 ± 0.6 % in ISO-stimulated cells (*P* = 0.008). These data suggest that ISO activation of *βAR*s, stimulates movement of Ca_V_1.2 out of EEs and into another cellular compartment.

**Figure 1.**
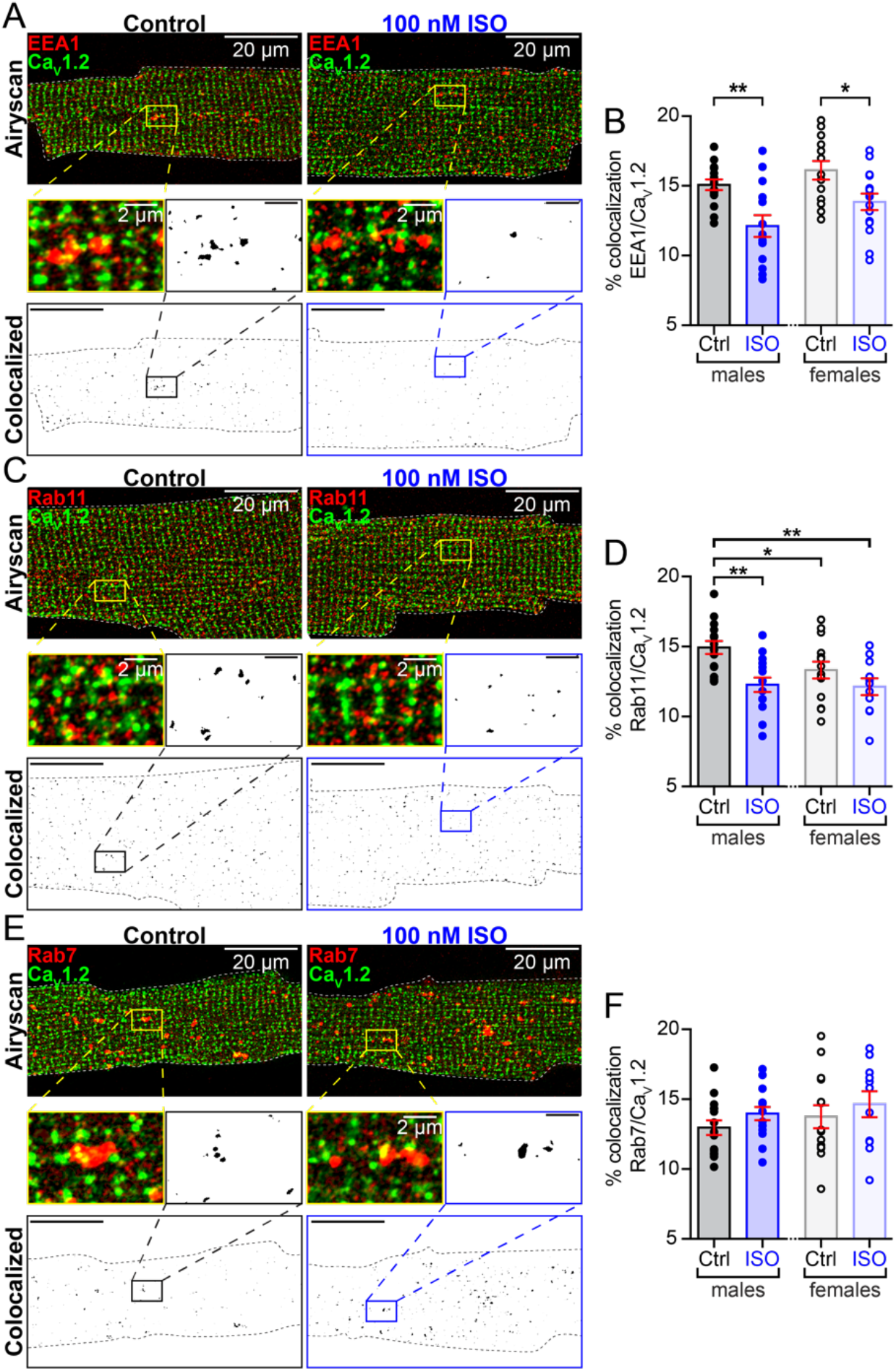
Internal pools of presynthesized Ca_V_1.2 channels reside on endosomes. (*A*) Two-color Airyscan super-resolution images of control and 100 nM ISO-stimulated, AMVMs immunostained to examine distributions of Ca_V_1.2 and EEA1 positive early endosomes. Binary colocalization maps (*bottom*) display pixels in which Ca_V_1.2 and endosomal expression precisely overlapped. (*B*) Histograms showing % colocalization between EEA1 and Ca_V_1.2 in male and female AMVMs, in control (male: *N* = 3, *n* = 15; female *N* = 3, *n* = 13) and ISO-stimulated conditions (male: *N* = 3, *n* = 14; female *N* = 3, *n* = 15). (*C*) Immunostaining of Ca_V_1.2 and Rab11 positive recycling endosomes and (*D*) accompanying histogram summarizing results from control (male: *N* = 4, *n* = 15; female *N* = 3, *n* = 14) and ISO conditions (male: *N* = 3, *n* = 15; female *N* = 3, *n* = 11). (*E*) Immunostaining of Ca_V_1.2 and Rab7 positive LEs and lysosomes. (*F*) Histogram summarizing results from control (males: *N* = 3, *n* = 15, females: N = 3, n = 14) and ISO-stimulated AMVMs (males: *N* = 3, *n* = 15, females: N = 3, n = 11). Error bars indicate SEM. Two-way ANOVA **P < 0.01; *P < 0.05. Pictured are male AMVMs, for females see Figure S1.

Cargo exiting the EE can be routed either to REs, LEs, or back to the sarcolemma via the fast recycling pathway (see working model in Figure 7A) (17, 19). If *βAR* activation stimulates Ca_V_1.2 channel trafficking from EEs into REs then a testable prediction is that ISO should increase colocalization between Ca_V_1.2 and Rab11 (a marker of REs). Accordingly, we examined the distribution of Ca_V_1.2 on Rab11-positive REs, and found a population of Ca_V_1.2-cargo-carrying REs, where Ca_V_1.2 colocalized with 14.9 ± 0.5 % of Rab11-positive pixels in males and with 13.3 ± 0.6 % in females (Figure 1C and D). An 18% decrease in colocalization between Ca_V_1.2 and Rab11 was observed in male cells treated with 100 nM ISO (*P* = 0.002). Data from female cardiomyocytes showed a similar downward trend in colocalization. These results do not support the hypothesis that Ca_V_1.2 channels move from Rab4-positive EEs into REs in response to ISO. We next tested the hypothesis that ISO stimulation drives the trafficking of Ca_V_1.2 channels from EEs to LEs, however despite identifying a population of Ca_V_1.2 channels on Rab7-positive LEs in male and female ventricular myocytes, ISO stimulation did not significantly alter the degree of Rab7/Ca_V_1.2 colocalization (Figure 1E and F; *P* = 0.28 in males, *P* = 0.39 in females). Having ruled out two of the three possibilities, we surmise that *β*AR stimulation drives an intracellular pool of Ca_V_1.2 channels from Rab4-positive EEs into the fast recycling pathway, and Rab11-positive REs into the slow recycling pathway.

### ISO-stimulated enhancement of Ca_V_1.2 recycling is regulated by Rab4 and Rab11

Having determined that *βAR* stimulation decreases the number of Ca_V_1.2 channels on EEs and REs we next tested if Rab4-dependent fast and, Rab11-dependent slow recycling pathways facilitate enhancement of Ca_V_1.2 delivery into the PM of transiently transfected tsA201 cells. While these cells do not recapitulate all of the intricacies of signaling in cardiomyocytes, we utilized them here because, 1) they provide a reductionist framework on which to test our Rab4 and Rab11 hypotheses in the absence of other voltage-gated channels; 2) they can be easily transfected, permitting manipulation of Rab-protein complement; and 3) they endogenously express *βARs* (24). PM expression of Ca_V_1.2 channels was monitored during activation of *βARs* with 100 nM ISO. Clusters of channels were readily identified in the TIRF footprint of the cell (Figure 2A). Upon wash-in of ISO, Ca_V_1.2-tagRFP intensity in the TIRF footprint increased by an average of 1.37-fold over a period of minutes (Figure 2B; τ_on_ = 2.26 ± 0.02 min), in close agreement with previous results showing a 44.9 ± 6.2 % increase in channels in ISO-stimulated AMVM sarcolemmas (23). An increased density (number/μm^2^) of channel clusters in the TIRF footprint contributed to this elevation (Figure 2C; *P* < 0.0001). These results suggest that ISO stimulates enhanced recycling of Ca_V_1.2 channels to the PM.

**Figure 2.**
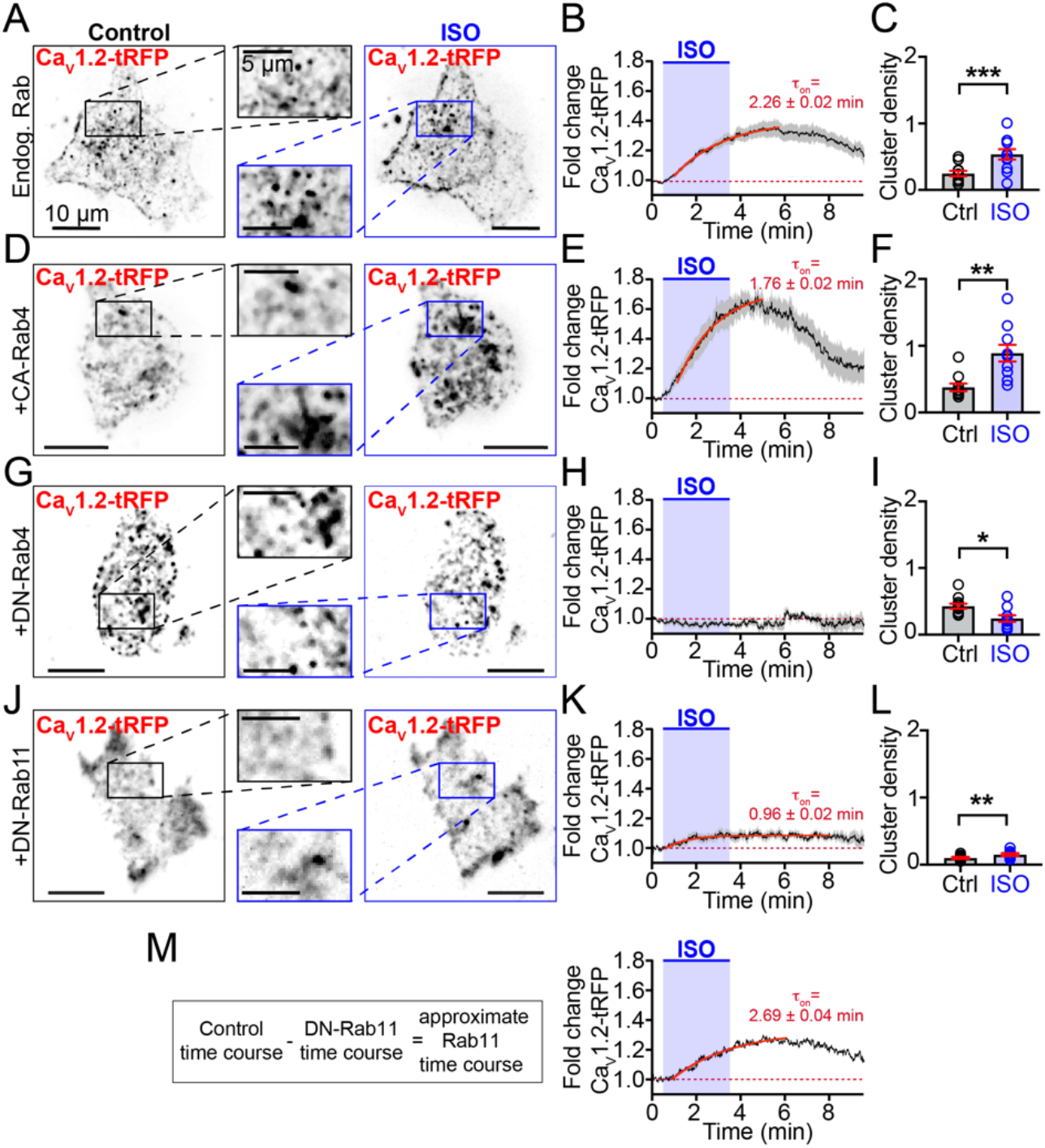
Rab4a and Rab11a regulate an ISO-stimulated boost in Ca_V_1.2 recycling. (*A*) TIRF images of Ca_V_1.2-tRFP distribution in the PM of tsA-201 cells before (*left*) and after 100 nM ISO (*right*; *n* = 18). (*B*) Time course and kinetics (solid red line) of the fold change in Ca_V_1.2-tRFP intensity in the TIRF footprint before, during and after ISO stimulation. (*C*) Histogram summarizing Ca_V_1.2 channel cluster density (number/μm^2^) in the TIRF footprint in control and ISO stimulated conditions. (*D-F*) Same layout format in tsA-201 cells co-expressing constitutively active (GTP-locked) Rab4a (*n* = 10). (*G-I*) Same layout format in tsA-201 cells co-expressing dominant negative (GDP-locked) Rab4a (*n* = 12). (*J-L*) Same layout format in tsA-201 cells co-expressing dominant negative (GDP-locked) Rab11a (*n* = 8). (*M*) Calculation and resultant theoretical time-course and kinetics of Rab11a-dependent ISO-stimulated recycling.

We tested the role of Rab4a in this dynamic response by co-expressing a mutant Rab4a with a single amino acid substitution (Q67L) that renders it resistant to GTP-hydrolysis, locking it in a GTP-bound, constitutively active state (CA-Rab4a^Q67L^; Figure 2D-F) (25). In these cells, addition of ISO stimulated a 1.69-fold increase in intensity, almost twice the maximal response observed in controls, and increased cluster density 2.4-fold (Figure 2F; *P* = 0.002). Interestingly, co-expression of CA-Rab4a had no effect on basal Ca_V_1.2 cluster density before addition of ISO (*P* = 0.08; Figure 2C and F) indicating that even in this GTP-locked active state, ISO-stimulation is required to trigger the boost in Ca_V_1.2 recycling. After ISO, the GTP-bound CA-Rab4a facilitates a larger, faster (1.76 ± 0.02 min) recycling response. This suggests that Rab4a plays a role in *β*AR-stimulated channel recycling but also implies the involvement of an upstream effector.

To support this postulate, we examined the ISO response in cells expressing a dominant-negative, GDP-locked variant of Rab4 (DN-Rab4^S22N^). Under these conditions, ISO-application failed to enhance Ca_V_1.2 surface expression and instead a slight decrease in Ca_V_1.2-tagRFP intensity and cluster density was observed in the PM over the course of the experiment (Figure 2G, H, and I). These results confirm that Rab4a is part of the essential trafficking machinery that underlies the ISO-stimulated enhanced recycling of Ca_V_1.2.

Since a subpopulation of Ca_V_1.2 channels that localize to Rab11 positive REs was also identified in AMVMs (Figure 1C and D), we tested the role of Rab11a in this dynamic recycling response to ISO using a dominant negative, GDP-locked DN-Rab11a^S25N^. Despite impaired Rab11a function, an ISO-stimulated increase in Ca_V_1.2-tagRFP intensity was still evident, albeit to a lesser extent than in controls (1.12-fold; Figure 2J and K). This was accompanied by a 1.49-fold increase in cluster density (*P* = 0.009; Figure 2L). Accordingly, DN-Rab11a-mediated knock down of Rab11a activity generated ~33% of the response seen in controls while knock down of Rab4a abolished the response (Figure 2H and M). These data suggest that upstream Rab4a activity is necessary, not only for fast recycling to the PM but also for transfer of Ca_V_1.2 cargo from EEs to REs. Impaired Rab4a function creates a ‘road block’ in the endosomal recycling system. The role of Rab4a in fast recycling was unmasked in cells with knocked down Rab11a activity, where τ_on_ of the stimulated recycling response was 0.96 ± 0.02 min (Figure 2K), significantly faster than controls where both Rab4a and Rab11a were active (2.26 ± 0.02 min). A theoretical time course of the slow Rab11a-dependent contribution was calculated by subtracting the predominantly Rab4a-mediated recycling time-course in DN-Rab11a expressing cells from endogenous Rab controls (Figure 2M). This theoretical Rab11a-dependent response displayed a 1.29-fold increase in Ca_V_1.2-tagRFP intensity and was well-fit with a single exponential function with a τ_on_ = 2.69 ± 0.04 min that was slower than control or Rab4a-mediated responses. Based on these data, our results suggest that Ca_V_1.2 channels stimulated to recycle to the PM in response to *βAR* activation are sourced approximately 1/3^rd^ from the fast Rab4a, and 2/3^rd^ from the slow Rab11a recycling pathways.

### Actin and microtubule disruption impairs ISO-stimulated Ca_V_1.2 recycling

We used Airyscan super-resolution microscopy to examine Ca_V_1.2 channel proximity to microtubules (MTs) and actin filaments (Figure 3A-B). Immunostaining with α-tubulin revealed the extensive MT cytoskeleton with its lattice, grid-like appearance in the subsarcolemma, transitioning to a more longitudinally-oriented network deeper in the cell interior (Figure 3A). Ca_V_1.2 channels decorate the MTs in both locations. Phalloidin-staining of the actin network revealed the periodic alignment of sarcomeric actin (Figure 3B). It is notoriously difficult to visualize cortical actin in cardiomyocytes due to the overwhelming abundance of sarcomeric actin but in many locations Ca_V_1.2 channels were co-localized with actin (Figure 3B, white arrowheads). The degree of colocalization between Ca_V_1.2 and, in particular MTs, implies a role for the cytoskeleton in regulating channel availability.

**Figure 3.**
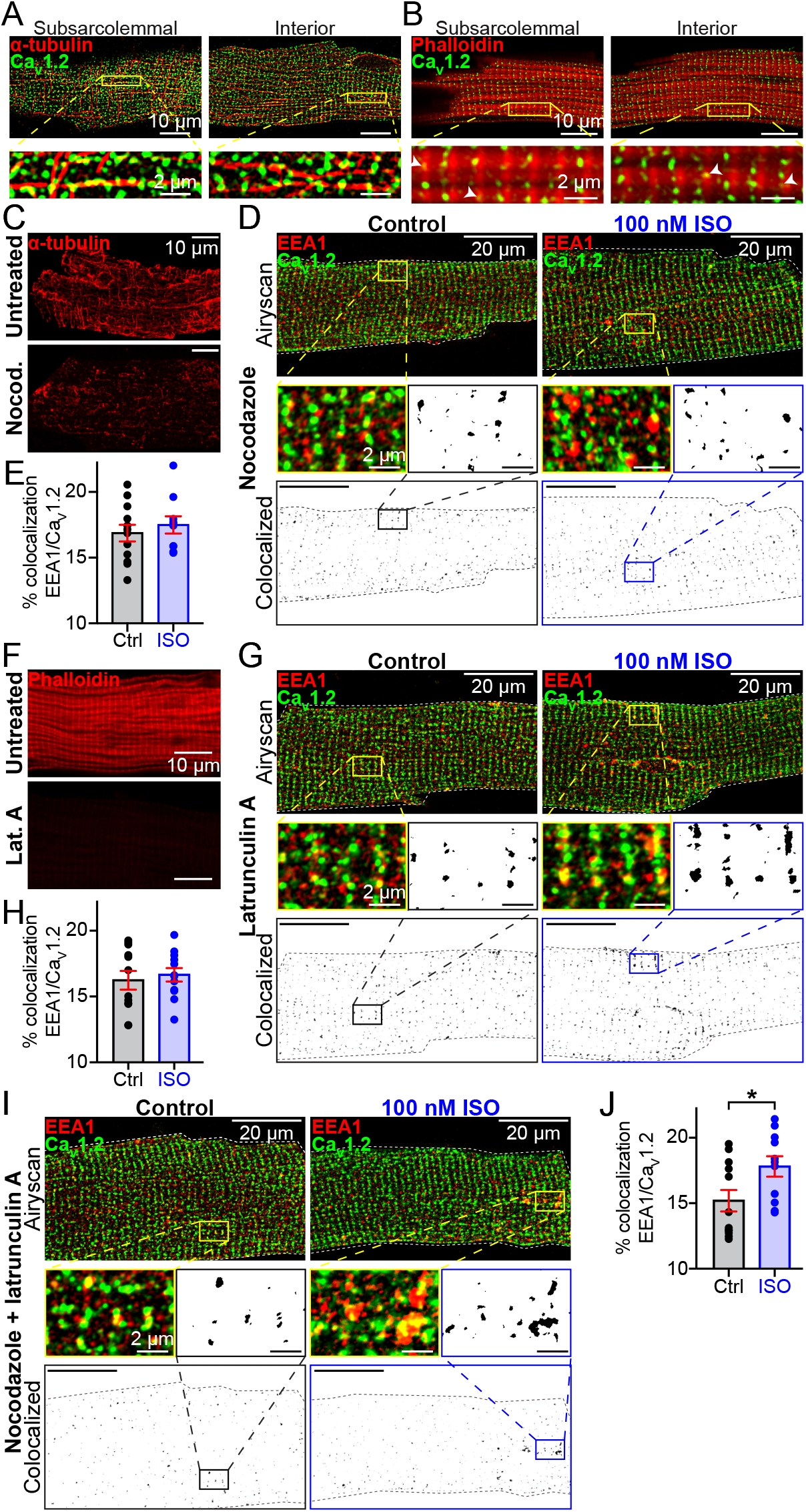
Actin and MT polymerization are essential for ISO-stimulated Ca_V_1.2 recycling. Two-color Airyscan images of fixed AMVMs immunostained to examine the relative localization of Ca_V_1.2 and (*A*) α-tubulin, or (*B*) phalloidin-stained actin. (*C*) shows the distribution of α-tubulin in untreated (*top*) and nocodazole-treated AMVMs (*bottom*). (*D*) Airyscan images of Ca_V_1.2 and EEA1 distribution in nocodazole-treated control (*N* = 3, *n* = 12; *left*) and ISO-stimulated (*N* = 3, *n* = 10; *right*) AMVMs. *Bottom*: binary colocalization maps display pixels in which Ca_V_1.2 and EEA1 expression precisely overlapped. (*E*) Histogram summarizing % colocalization of EEA1 with Ca_V_1.2 in nocodazole-treated cells. (*F*) Actin distribution in untreated (*top*) and latrunculin-A treated cells (*bottom*). (*G*) Airyscan images and binary colocalization maps of Ca_V_1.2 and EEA1 distribution in latrunculin-A-treated control (*N* = 3, *n* = 12; *left*) and ISO-stimulated (*N* = 3, *n* = 12; *right*) AMVMs. (*H*) Histogram showing % colocalization of EEA1 with Ca_V_1.2 in latrunculin-A-treated AMVMs. (*I*) Airyscan images and binary colocalization maps of AMVMs treated with both latrunculin-A and nocodazole under control (*N* = 3, *n* = 12) and ISO stimulated conditions (N = 3, n = 11), with accompanying summary histogram (*J*).

To study the role of these cytoskeletal highways in *βAR*-stimulated Ca_V_1.2 mobilization from endosomes, we examined EEA1-localized Ca_V_1.2 channel populations in AMVMs treated with cytoskeletal disruptors. Accordingly, freshly isolated myocytes were treated for 2 hrs with 10 μM nocodazole, a drug known to prevent addition of tubulin to dynamic MTs, and depolymerize the stable variety (26). Immuno-staining with anti-α-tubulin confirmed the treatment had substantially disordered the MTs (Figure 3C). In this and upcoming experimental series, we refer to cells that did not receive cytoskeletal disruptors as ‘untreated’. As in untreated cells (Figure 1A and B), Ca_V_1.2 was observed to colocalize with a sub-population of EEs (16.9 ± 0.6 %; Figure 3D and E). However, in cells stimulated with 100 nM ISO prior to fixation, the reduction in colocalization between EEA1 and Ca_V_1.2 we had previously observed in untreated cells was absent in nocodazole-treated cells, instead remaining at 17.5 ± 0.7 % (*P* = 0.51 compared to nocodazole-treated control). These data suggest that MT network disruption prevents the ‘emptying’ of Ca_V_1.2 channel cargo from the EEs into the sarcolemma and support a role for MTs and their associated motor proteins as conduits for ISO-stimulated Ca_V_1.2 recycling.

We then examined the role of the actin cytoskeleton by disrupting it with latrunculin A (lat-A; 5 μM for 2 hrs), which both facilitates F-actin depolymerization (27) and prevents polymerization by sequestering actin monomers (26). Importantly, lat-A specifically affects the actin cytoskeleton but spares the MT network (28). Alexa Fluor 647-conjugated phalloidin staining of filamentous-actin (F-actin) was used to visually confirm lat-A mediated actin-disruption (Figure 3F). In lat-A-treated myocytes, Ca_V_1.2 and EEA1 colocalization was similar to untreated controls (16.2 ± 0.7 % versus 15.1 ± 0.4 %; *P* = 0.14). However, ISO-stimulation did not affect colocalization levels (16.7 ± 0.5 %; *P* = 0.64; Figure 3G and H). In addition, in cells co-treated with nocodazole and lat-A, ISO-stimulation actually promoted a small, but significant increase in colocalization between Ca_V_1.2 and EEA1 (*P* = 0.03; Figure 3I and J). These data indicate a profound alteration in the endosomal pathway where recycling and or endocytosis have been impaired, creating an ‘endosomal traffic jam’ leading to accumulation of cargo on the endosomes, with no cytoskeletal highways to transport the cargo to its destination.

### Cytoskeletal disruption alters ISO-stimulated Ca_V_1.2 dynamics and recycling

Real-time visualization and quantification of the effects of cytoskeletal disruption on channel trafficking was performed using transduced AMVMs isolated from mice that had received a retro-orbital injection of AAV9-Ca_V_β_2a_-paGFP. This auxiliary subunit of Ca_V_1.2 binds to the pore-forming subunit with a 1:1 stoichiometry and acts in this context as a biosensor reporting the location of the subset of Ca_V_1.2 α_1c_ it interacts with. This approach was previously validated by our group (23), with super-resolution microscopy experiments confirming that the biosensor and the α_1c_ colocalize, and unlike overexpression of α_1c_, at these concentrations we have found that Ca_V_β_2a_-paGFP transduction does not appreciably affect Ca_V_1.2 α_1c_ expression as indicated by unaltered basal channel cluster sizes. A further advantage of this approach is that it requires no culturing of AMVMs which are known to rapidly dedifferentiate in culture (29). We began by examining the dynamic channel trafficking response to ISO in untreated AMVMs (i.e., in the absence of cytoskeletal disruptors) using TIRF microscopy. Discrete puncta of Ca_V_β_2a_-paGFP decorated the TIRF footprint of the myocyte during control frames and additional puncta/clusters were seen to appear in the TIRF footprint supplementing the initial complement after perfusion with 100 nM ISO (Movie 1 and Figure 4A). Given our endosome/Ca_V_1.2 immunostaining results (Figure 1), this may represent endosomal cargo mobilized in response to ISO from subsarcolemmal locations deeper within the cell. In agreement with this, 3D-plots of Ca_V_β_2a_-paGFP intensity over time and cell depth, constructed from 4D-spinning disk confocal experiments performed on transduced AMVMs at 37°C, indicated Ca_V_β_2a_-paGFP was mobilized from several microns within the cell, and moved toward the surface in response to ISO (Figure S3). At physiological temperature, following a delay of ~3 sec, the response to ISO proceeded monoexponentially with a τ = 2.53 ± 0.28 sec until Ca_V_β_2a_-paGFP intensity reached a plateau (Figure S3C-D), presumably achieved when the endosomal pool of channels had been depleted and balance between insertion and endocytosis reached a new equilibrium. These data fit well with previous measurements of the time course of I_Ca_ responses to ISO (30, 31). Interestingly, exchange of the perfusate with one containing 100 nM of angiotensin II, quickly reversed the response, and a subset of Ca_V_β_2a_-paGFP moved back into the interior of the cell with a τ = 2.94 ± 0.96 sec. Previous work performed at room temperature reported that prolonged (30 - 60 min) application of angiotensin II produces Ca_V_1.2 channel internalization (32), but here at physiological temperature, the time course was significantly accelerated.

**Figure 4.**
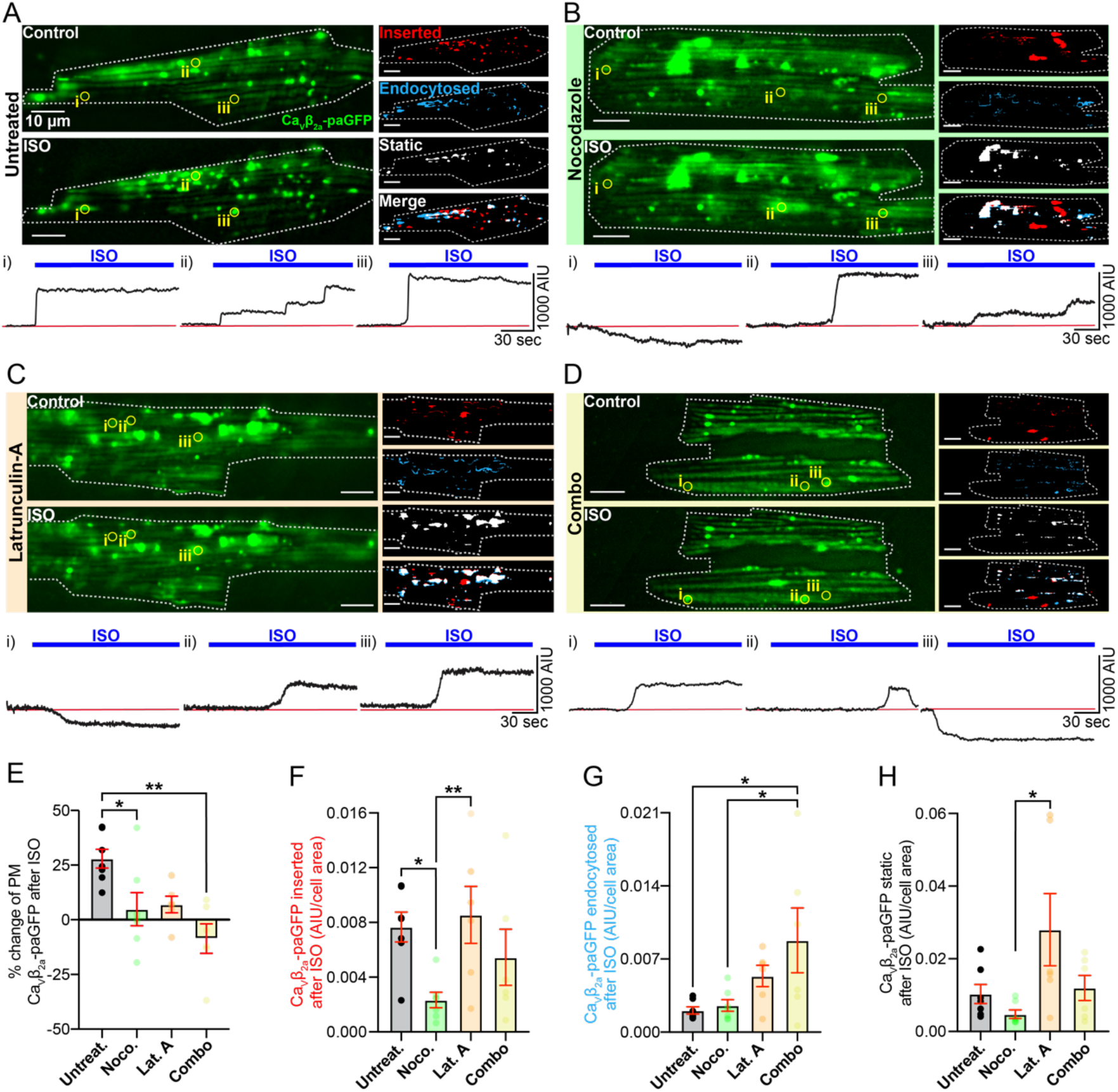
Dynamic imaging to unmask the mobile channel population. (*A*) TIRF images of GFP fluorescence emission from Ca_v_β_2a_-paGFP transduced AMVMs before (*top*) and after 100 nM ISO (*bottom*). Images illustrating inserted, endocytosed, static, and merged channel populations are shown to the right. *Bottom*: Time course of the changes in Ca_v_β_2a_-paGFP intensity in ROIs (i-iii) indicated by yellow circles on TIRF images. (*N* = 5, *n* = 7). Same format for cells pre-treated with (*B*) nocodazole (*N* = 5, *n* = 7), (*C*) latrunculin-A (*N* = 3, *n* = 6), or (*D*) both cytoskeleton disruptors (*N* = 3, *n* = 6). (*E-H*) Histograms summarizing statistics for these experiments.

To test the hypotheses that ISO treatment increases sarcolemmal expression of Ca_V_1.2 by stimulating channel insertion/recycling, and that cytoskeletal highways carry these recycling channels to their destination, we performed ‘image math’ (see Methods) on TIRF time series to quantify the subpopulations of Ca_V_β_2a_-paGFP in the TIRF-footprint that were: i) inserted, ii) endocytosed, and iii) stably expressed during ISO stimulation. Responses to ISO in untreated AMVMs were compared, to those in cells treated with nocodazole, lat-A, or a combination of both (Figure 4). Live cell time series experiments revealed a dynamic population of Ca_V_β_2a_-paGFP in all cells examined (see Movies 1-4), although the dynamics were appreciably less in cells that received cytoskeletal disrupting treatments, supporting the idea that both F-actin and MTs are important conduits of this response. The number of channels at the sarcolemma at any given time, is dictated by the balance between channel insertions via the biosynthetic delivery and endosomal recycling pathways, and channel removals via endocytosis. In untreated cells, ISO stimulation heavily shifted the balance in favor of insertion, implying a stimulated insertion/recycling process (Figure 4A, and E-G). This mismatch between insertion and endocytosis produced a 27.95 ± 4.31 % increase in sarcolemmal Ca_V_β_2a_-paGFP expression (Figure 4E). The time course of ISO-stimulated insertions of channels in untreated AMVMs can be observed in the ROIs highlighted in Figure 4A. Examination of these time courses revealed rapid step-like insertion profiles, suggesting that Ca_V_1.2 appear to often insert into the sarcolemma as preformed clusters, containing many channels (Figure 4A(i-iii), B(ii and iii), C(ii and iii), D(i and ii)). In some cases, a delivery hub was evident, where a succession of channel clusters appeared to insert one after the other (Figure 4A(ii) and B(iii)). Channel endocytosis also appeared to involve removal of channel clusters in some instances (Figure 4C(i), D(iii)), while in other ROIs, a slower, gradual removal of potentially individual channels was more evident (Figure 4B(i)).

Recycling of endosomal cargo back to the PM is known to rely on both MTs and actin (33). Indeed, cytoskeletal disruption reduced the magnitude of the ISO-stimulated augmentation of sarcolemmal Ca_V_β_2a_-paGFP expression compared to untreated cells (Figure 4E). Interestingly, how the two elements of the cytoskeleton affected the ISO-stimulated change in sarcolemmal expression varied somewhat, as revealed by examination of insertion and endocytosis events in each AMVM cohort. Anterograde transport and targeting of Ca_V_1.2 to the t-tubule membrane is known to occur along MTs, anchored there via the BAR-domain containing protein, BIN1 (A.K.A. amphiphysin II) (34). Here, our experiments were focused not on long distance trafficking from the trans-golgi to the membrane, but rather from the local endosome pool of channels and thus channel dynamics were observed over short 3 min periods before and after application of ISO. Our data indicates that nocodazole mediated MT disruption reduced ISO-stimulated insertion of Ca_V_β_2a_-paGFP by an average of ~70% compared to untreated cells (Figure 4F). Channel internalization and the lifetime of channels in the membrane was not affected by the degree of MT disruption tested here, as both endocytosed and static channel population sizes were not significantly altered by this treatment (Figure 4G and H).

Actin disruption with lat-A in contrast had no significant effect on channel insertion compared to untreated cells (Figure 4F). In neurons and HEK293 cells, interactions between Ca_V_1.2 and the actin-interacting protein α-actinin have been found to stabilize Ca_V_1.2 channel expression at the PM (35, 36). A notable trend toward increased endocytosis of Ca_V_β_2a_-paGFP in lat-A treated cells was detected but this failed to reach significance assessed by a one-way ANOVA test (Figure 4G). There was also a trending increase in the fraction of stable Ca_V_β_2a_-paGFP in the TIRF footprint that appeared to be left ‘stranded’ there (Figure 4H), perhaps reflective of incomplete internalization of Ca_V_β_2a_-paGFP due to the lack of actin dynamics and the disrupted cortical actin network. This effect was not statistically different from controls but was significantly different from nocodazole treated cells. Finally, combined treatment with nocodazole and lat-A actually *reduced* the overall expression of Ca_V_β_2a_-paGFP in the TIRF footprint by 8.63 ± 6.76 % over the course of the experiment (Figure 4F and G). This occurred when the balance of endocytosis and insertion/recycling shifted in favor of endocytosis. Collectively, these data suggest that MTs and actin are both important conduits of *βAR*-stimulated Ca_V_1.2 recycling, with MTs playing the major role in channel insertion.

### βAR mediated I_Ca_ regulation is abrogated by cytoskeletal disruption

To ascertain whether ISO-stimulated recycling of Ca_V_1.2 channels into the cardiomyocyte sarcolemma makes any functional contribution to the *βAR* regulation of these channels, we performed whole cell patch clamp recordings on freshly isolated AMVMs under various cytoskeletal disrupting conditions. Stimulation of untreated cardiomyocytes with 100 nM ISO, produced a significant 1.65 ± 0.19-fold enhancement of *I*_Ca_ (*P* = 0.04; Figure 5A and B) and caused a 13.33 ± 1.49 mV leftward-shift in the voltage-dependence of conductance (measured as the difference between V_1/2_ of each fit; *P* < 0.0001; Figure 5C and Table 1). In addition, we observed an increase in the slope steepness of the Boltzmann function used to fit the G/G_max_ data from 5.45 ± 0.84 in controls to 4.90 ± 1.06 in 100 nM ISO (Figure 5C and Table 1), suggesting a potential increase in cooperative gating behavior (23).

**Figure 5.**
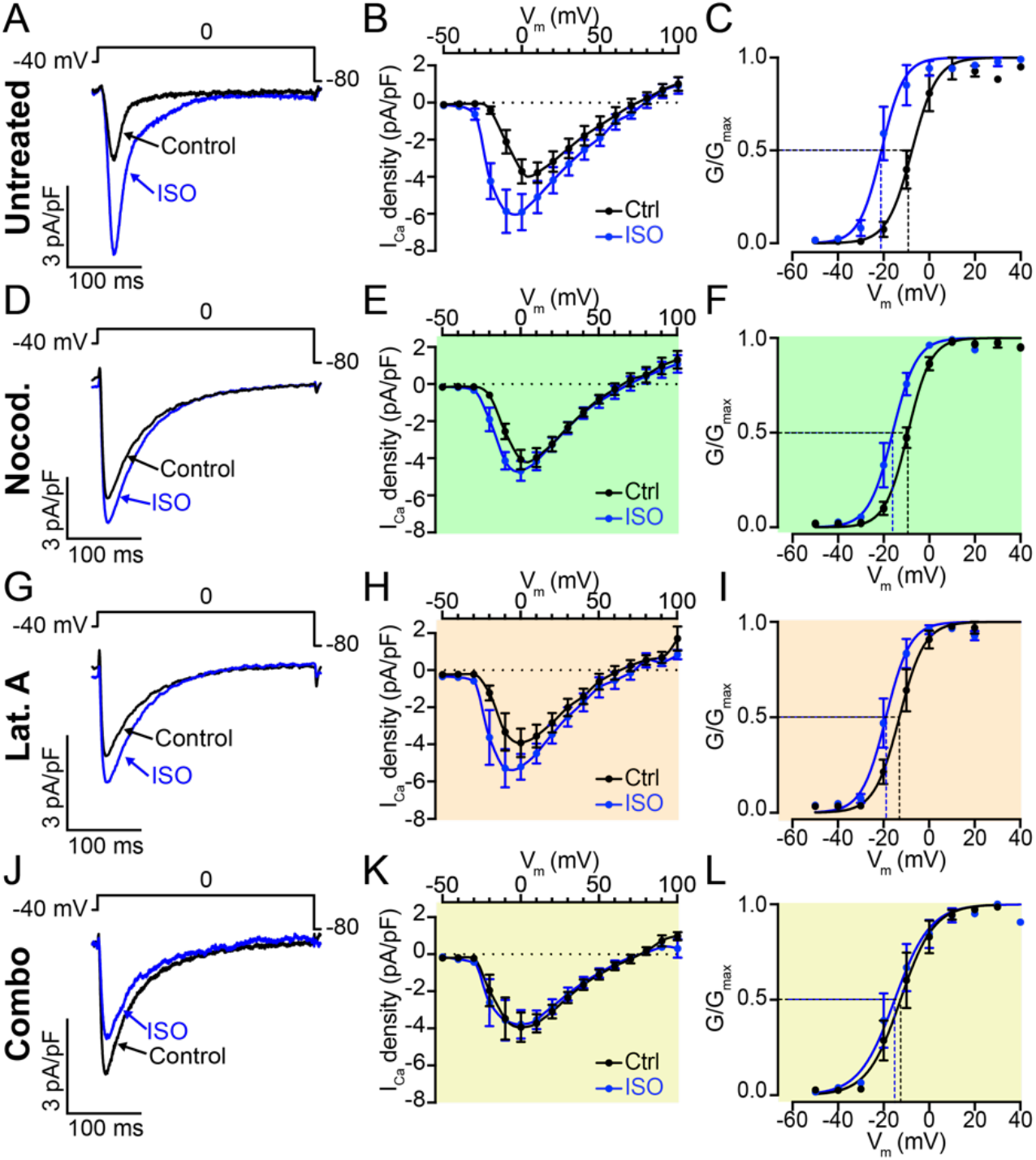
*β*AR mediated I_Ca_ enhancement is blunted by reducing dynamic channel insertion. (*A*) Whole cell currents elicited from a representative AMVM during a 300 ms depolarization step from −40 mV to 0 mV before (control: black) and during application of 100 nM ISO (blue). (*B*) I-V plot summarizing the results from *n* = 7 cells (from *N* = 7 animals) subjected to test potentials ranging from −50 mV to +100 mV. Currents were normalized to cell capacitance to generate current density. (*C*) Voltage-dependence of the normalized conductance (G/G_max_) before and during ISO application, fit with Boltzmann functions. (*D-F*) show whole cell currents (*D*), I-V (*E*), and G/G_max_ plots (*F*) from AMVMs pre-treated for 2 hrs with 10 μM nocodazole (*N* = 4, *n* = 7). (*G-H*) show the same for cells treated with 5 μM lat-A (*N* = 5, *n* = 5). (*J-L*) show results from cells treated with a combination of both (*N* = 3, *n* = 6).

**Table 1.**

| | Fold-change in peak $I_{Ca}$ | $V_{1/2}$ (mV) | | Slope factor | |
| --- | --- | --- | --- | --- | --- |
|  |  | Control | ISO | Control | ISO |
| **Untreated** | $1.65 \pm 0.19^*$ | $-7.57 \pm 0.95$ | $-20.90 \pm 1.14^{****}$ | $5.45 \pm 0.84$ | $4.90 \pm 1.06$ |
| **Nocodazole** | $1.18 \pm 0.08$ | $-9.38 \pm 0.44$ | $-16.10 \pm 0.84^{***}$ | $5.03 \pm 0.41$ | $5.27 \pm 0.72$ |
| **Latrunculin A** | $1.39 \pm 0.13$ | $-13.03 \pm 0.87$ | $-18.95 \pm 0.97^*$ | $5.51 \pm 0.76$ | $5.33 \pm 0.88$ |
| **Noco + Lat.A** | $1.01 \pm 0.11$ | $-12.62 \pm 1.40$ | $-15.16 \pm 1.53$ | $7.32 \pm 1.24$ | $7.82 \pm 1.36$ |

The impact of MT disruption on the *I*_Ca_ response to ISO was tested in AMVMs incubated with nocodazole. This dose and duration of treatment did not alter control *I*_Ca_ amplitude (compared to untreated control). Indeed, none of the cytoskeletal disrupting treatments had any significant effect on control *I*_Ca_ with all of them peaking between −3.78 and −4.07 pA/pF (Figure 5B, E, H, and K). However, nocodazole blunted the *I*_Ca_ response to ISO by ~72% (1.18 ± 0.08-fold increase versus 1.65-fold change in untreated cells), halved the magnitude of leftward-shift of the voltage dependence of conductance (6.73 ± 1.38 mV versus the 13.33 mV shift in untreated cells), and eliminated the tendency toward cooperativity indicated by the slope of the Boltzmann function used to fit the G/G_max_ (Figure 5D-F; Table 1).

We used the same experimental paradigm to test whether the actin cytoskeleton plays any role in the functional regulation of Ca_V_1.2 by *βAR*s. Cells treated with lat-A exhibited a 40% reduction in the I_Ca_ augmentation response to ISO compared to controls (Figure 4G and H, Table 1), and a similar halving of the leftward-shift in the voltage dependence of conductance as in nocodazole-treated cells (5.92 ± 1.63 shift; Figure 5I and Table 1). ISO stimulation produced G/G_max_ data that was well-fit with a Boltzmann function that was less steep than in ISO-stimulated untreated cells but still indicated enhanced cooperativity (Figure 5I and Table 1).

Finally, we examined the effect of ISO on *I*_Ca_ when both MTs and actin filaments were disrupted with a combined nocodazole and lat-A treatment. Under these conditions, *βAR*-mediated regulation of Ca_V_1.2 was essentially abolished (Figure 5J-K), with ISO, generating only a 1.01 ± 0.11-fold increase in *I*_Ca_, representing a 98% reduction in the response compared to untreated cells. The leftward-shift in the voltage dependence of conductance was reduced to 2.54 ± 1.49, equivalent to only 20% of the response seen in untreated cells. The slope factor of the Boltzmann-function used to fit the data was less steep than untreated cells in both control and ISO-stimulated conditions, indicating reduced cooperativity. These functional data collectively indicate that an intact cytoskeleton is an essential requirement for *βAR*-mediated regulation of Ca_V_1.2.

### Clustering of Ca_V_1.2 channels is supported by the cytoskeleton

The observation that channel cooperativity was altered in AMVMs with disrupted cytoskeletal elements implies that the cytoskeleton might be important not only for *βAR*-mediated regulation of Ca_V_1.2 but also for the stabilization and support of channel clusters. We tested this idea by examining Ca_V_1.2 channel distribution with <u>G</u>round <u>S</u>tate <u>D</u>epletion (GSD) super-resolution nanoscopy. ISO-stimulation resulted in the formation of Ca_V_1.2 channel super-clusters in AMVMs which were, on average, 22.6% larger than the clusters in control AMVMs (Figure 6A and E). Super-clustering could occur due to small clusters fusing together to form larger ones, or alternatively, could reflect enhanced sarcolemmal expression of Ca_V_1.2. If the superclusters form because of enhanced insertion/exocytosis of Ca_V_1.2 into the sarcolemma, then a testable prediction is that the intensity of the fluorescence emission (normalized to the cell area) should be increased. In contrast, if the super-clusters simply reflect fusion of existing sarcolemmal Ca_V_1.2 clusters then the total fluorescence intensity should be similar in control and ISO-treated cells. Accordingly, the normalized total integrated density was significantly larger in ISO-treated cells than controls (Figure 6F; *P* = 0.00002), in agreement with the idea that *βAR*-activation stimulates enhanced sarcolemmal insertion and resultant super-clustering of Ca_V_1.2 channels. Cytoskeletal disruption with nocodazole (Figure 6B), lat-A (Figure 6C), or a combination of the two (Figure 6D), did not affect basal channel expression in the sarcolemma as indicated by similar cluster areas and total integrated density values (Figure 6E-F). These results validate the unaltered *I_Ca_* observed in unstimulated cells from each of our four experimental groups (Figure 5) and supports the postulate that the lifetime of these channels in the membrane is longer than the 2 h cytoskeletal disruption period. In addition, ISO-stimulation failed to induce super-clustering or enhanced sarcolemmal expression of Ca_V_1.2 in AMVMs (Figure 6E-F). Altogether, these data suggest that both intact MTs and actin are necessary for the formation of Ca_V_1.2 super clusters in response to ISO stimulation.

**Figure 6.**
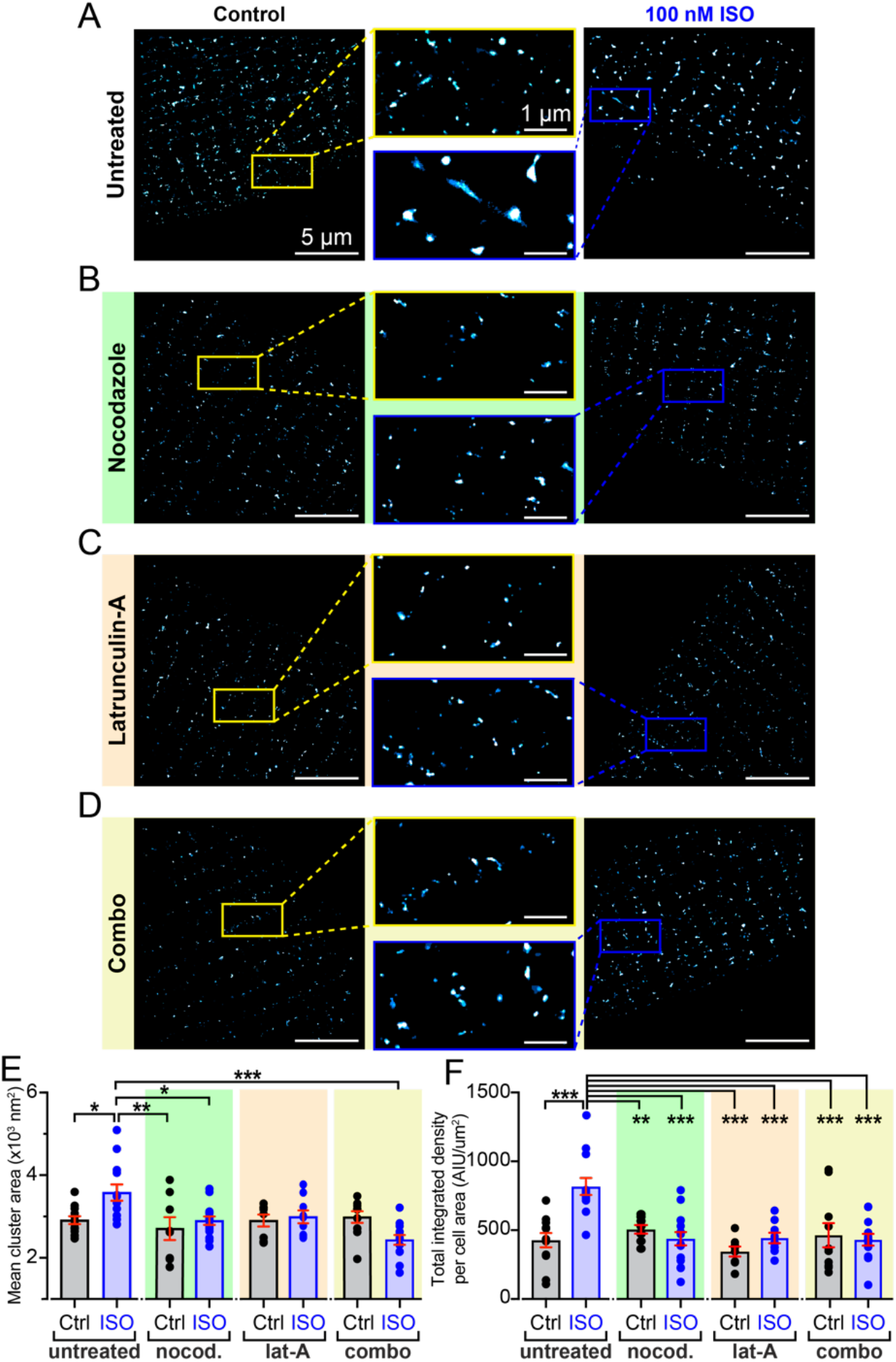
*β*AR-stimulated Ca_V_1.2 super-clustering and enhanced sarcolemmal expression requires an intact MT cytoskeleton. Super-resolution GSD localization maps of control (*left*) and 100 nm ISO-stimulated (*right*), fixed, AMVMs immunostained to examine Ca_V_1.2 channel distribution under (*A*) untreated (control: *N* = 3, *n* = 13; ISO: *N* = 3, *n* = 13), (*B*) nocodazole-treated (control: *N* = 3, *n* = 8; ISO: (*N* = 3, *n* = 15), (*C*) lat-A-treated (control: *N* = 3, *n* = 8; ISO: (*N* = 3, *n* = 9), and (*D*) combo-treated (control: *N* = 2, *n* = 10; ISO: (*N* = 3, *n* = 13) conditions. Maps were pseudocoloured ‘cyan hot’ and received a one-pixel median filter for display purposes. Yellow (*control*) and blue (*ISO*) boxes indicate the location of the zoomed-in regions displayed in the center. (*E*) and (*F*), aligned dot plots showing mean Ca_V_1.2 channel cluster areas and normalized total integrated density in each condition. Two-way ANOVA ***P<0.001, **P < 0.01, *P < 0.05.

## Discussion

The data presented in the present study provide the first report of an endosomal pool of Ca_V_1.2 channels in cardiomyocytes that undergoes rapid, targeted mobilization to the sarcolemma in response to *βAR*-activation, effectively creating an ‘on-demand’ trafficking pathway to facilitate a positive inotropic response during fight-or-flight. We present six major new findings: 1) intracellular pools of Ca_V_1.2 channels are present on EEs, REs, and in LEs and lysosomes in AMVMs; 2) *βAR*-activation triggers Ca_V_1.2 mobilization from EEs and REs to the sarcolemma via Rab4a-dependent fast, and Rab11a-dependent slow recycling pathways; 3) Ca_V_1.2 are often inserted or removed from the sarcolemma as large multi-channel clusters, rather than individual channels; 4) stimulated insertion of Ca_V_1.2 channels occurs along MTs; 5) the endosomal pool of Ca_V_1.2 is fueled by actin-dependent endocytosis; and finally, 6) adrenergic regulation of Ca_V_1.2 is abrogated by cytoskeletal disruption and loss of this dynamic recycling response. On the basis of these data, we present a new working model (Figure 7) for *βAR*-regulation of cardiac Ca_V_1.2 channels in which stimulated recycling of Ca_V_1.2, from sub-sarcolemmal pools of Rab4a and Rab11a-positive endosomes, results in enhanced expression of Ca_V_1.2 at the membrane of AMVMs. Resultant super-clustering and cooperative gating of Ca_V_1.2 channels contributes to the enhanced *I*_Ca_ and inotropic response.

**Figure 7.**
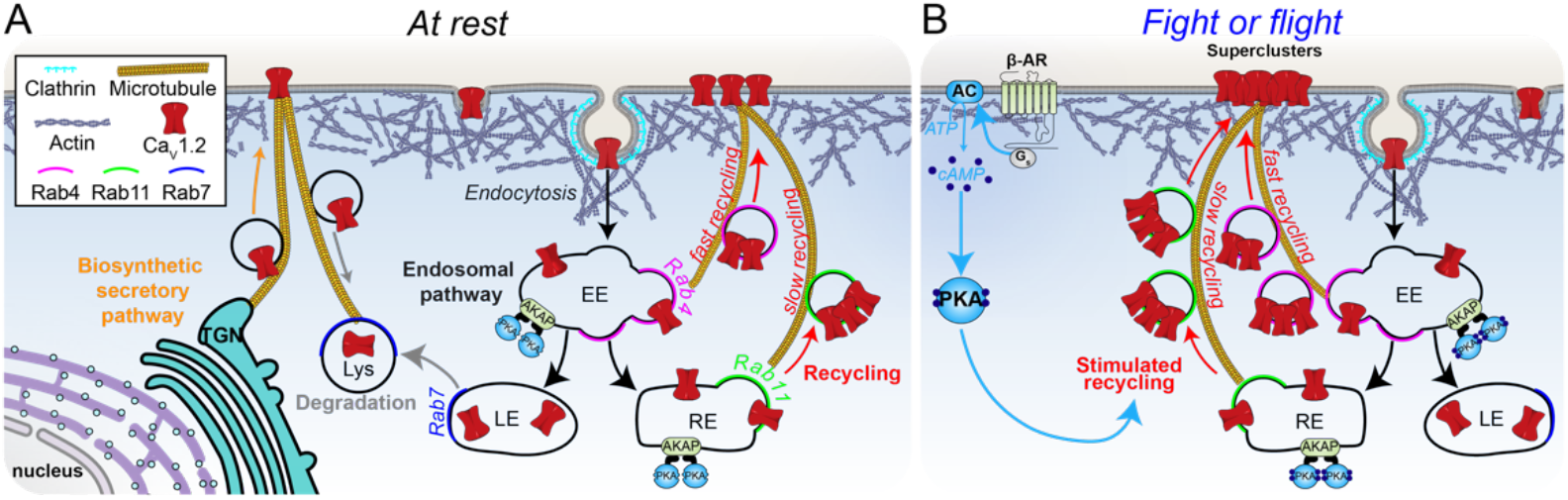
Our working model. *A*: Ca_V_1.2 channels on the ventricular myocyte sarcolemma are subject to dynamic equilibrium under resting, unstimulated conditions. Channels synthesized in the Golgi are transferred to negative-end anchored MTs at the trans-Golgi network (TGN) and transported in motor-protein shuttled vesicles, to plus-end anchored BIN1 hubs at the membrane. Steady state channel expression is achieved as ‘new’ channel insertions are balanced by ongoing channel endocytosis and recycling through the endocytic pathway. *B:* Activation of *βAR*s by ISO or during fight or flight, leads to stimulated recycling of Ca_V_1.2, from sub-sarcolemmal pools of Rab4a and Rab11a-positive endosomes. D-AKAP2 anchors PKA on endosomal membranes, future studies should investigate its role in *βAR*-stimulated channel recycling. Resultant ‘on-demand’ insertion of the endosomal reservoir of Ca_V_1.2, leads to augmented Ca_V_1.2 expression and super-clustering at the sarolemma, facilitates channel cooperativity, amplifies Ca^2+^influx, and contributes to the enhanced *I*_Ca_ and inotropic response.

Our data revealed pools of intracellular Ca_V_1.2 channels on EEA1 and Rab4-positive EEs, on Rab11 positive REs, and in Rab7 positive LEs and lysosomes. In response to ISO, EE and RE-localized channels underwent rapid recycling into the sarcolemma via the Rab4a-dependent fast recycling pathway and the slower, Rab11a recycling pathway. While this is the first report of stimulated recycling of a recruitable intracellular reservoir of Ca_V_1.2 in cardiomyocytes, small GTPase choreographed-recycling of endosome-localized ion channel pools are well-known to play a role in fine-tuning cellular responses to various stimuli including *βAR* stimulation. For example, in neurons, intracellular AMPA receptors (AMPAR) located on REs undergo Rab11-dependent recycling to the PM of dendritic spines in response to PKA-mediated phosphorylation of their GluA1 subunit at S845 downstream of *β2AR* stimulation (reviewed in (37, 38)). In the collecting ducts of the kidney, vasopressin release initiates a Gs-coupled signaling cascade that triggers PKA-mediated phosphorylation of aquaporin-2 (AQP2) at S256 and consequent recycling of AQP2 from Rab11-positive REs to the apical PM (39, 40). In the heart, acute stress initiates Rab11-dependent mobilization of endosomal reservoirs of SUR2-containing KATP channels and of KCNQ1-containing REs to the sarcolemma (41–44). Similarly, here we report an endosomal pool of Ca_V_1.2 channels that undergoes ‘on-demand’, stimulated recycling upon activation of *βAR*s, providing a functional reserve that drives ventricular inotropy during sympathetic stimulation.

Despite their fundamental importance in cardiac EC-coupling, the difficulty in visualizing and measuring Ca_V_1.2 channel dynamics means there is a remarkable lack of information about Ca_V_1.2 channel trafficking, endocytosis, and recycling in cardiomyocytes, with much of the published data focusing instead on data collected from heterologous expression systems. A previous study found that shRNA knockdown of Rab11a did not affect Ba^2+^ currents through Ca_V_1.2 in HEK293 cells or neonatal mouse ventricular myocytes while Rab11b knockdown significantly increased *IBa* (21), suggesting that Rab11b plays a contrary role to its close family member Rab11a, and actually regulates constitutive Ca_V_1.2 channel degradation. While we did not measure currents from tsA201 cells, it is reasonable to expect that currents should scale proportionally with the number of Ca_V_1.2 channels in the membrane, since the whole cell current (*I_Ca_*) is the product of the number of channels (*N*), open probability (*P_o_*), and single-channel current (*i_Ca_*), i.e., *I_Ca_ = N•i_Ca_•P_o_*. We found no evidence that knockdown of Rab11a function affected basal channel cluster density (*P* = 0.12) or number of channels in the membrane (measured with normalized mean grey values; *P* = 0.16) (compared to same-day controls with no Rab overexpression), in agreement with the findings of Kamp et al. However, knockdown of Rab11a function reduced the degree of *βAR*-stimulated Ca_V_1.2 insertion by 67%. This confirms the role of Rab11a in *βAR*-stimulated channel insertion rather than degradation.

Our work on live, AAV9-Ca_V_β_2a_-transduced AMVMs provides intriguing new insights into Ca_V_1.2 channel trafficking, and captures the complex dynamics of these channels. We find that these channels are often inserted into the sarcolemma as entire pre-formed clusters at nucleation sites. Sometimes, repetitive insertions were seen to occur at an individual site, conjuring an image of Ca_V_1.2-carrying endosomes queued up along MTs, anchored at a sarcolemmal delivery hub. Furthermore, channel endocytosis often appeared to occur via removal of entire clusters, while in other cases, a slower, gradual removal of channels suggested ongoing removal of individual channels. Our results at physiological temperature indicate that these insertion and removal events occur extremely rapidly and can be stimulated by activation of GPCR-signaling pathways. Activation of *βAR*s was observed to increase the probability of channel insertion while activation of AngII receptors increased the probability of channel removal. These very scenarios were predicted in a recently published computer model designed to test the hypothesis that ion channel clustering occurs via a stochastic self-assembly process (45). Our data provides answers to the hypotheticals raised by that model. Informing that models parameters with the experimental data acquired in this study would be an interesting sequel.

One well-characterized facet of cardiac Ca_V_1.2 channel trafficking is their targeted anterograde-delivery to the t-tubule membrane along BIN1-anchored MTs via the biosynthetic delivery pathway (34). Reduced levels of BIN1 in cardiomyocytes isolated from failing human hearts are associated with impaired Ca_V_1.2 channel delivery and slower onset calcium transients (10). Here, we find that Ca_V_1.2 channel recycling also occurs along MTs. Three independent lines of evidence support this conclusion. Firstly, MT disruption prevented ISO-stimulated mobilization of Ca_V_1.2 from EE pools (Figure 3D). Secondly, MT disruption significantly reduced stimulated channel insertions in AAV9-Ca_V_β_2a_-paGFP transduced AMVMs (Figure 4F and Movie 2). Thirdly, ISO-stimulated Ca_V_1.2 super-clustering was absent in nocodazole-treated AMVMs (Figure 6B and E). Our findings that basal Ca_V_1.2 channel distribution and *I_Ca_* amplitude were unaffected by 2 hr nocodazole treatment suggests that channel lifetime in the membrane is longer than 2 hrs, so that channels delivered along intact MTs prior to depolymerization by the drug, still largely remained there (Figure 6B). This agrees with previous measurements of PM Ca_V_1.2 channel half-times of ~3 hrs, although this has never been directly measured in cardiomyocytes (12). However, despite negligible effects on basal channel expression and function, inhibition of MT polymerization significantly reduced ISO-stimulated responses, blunting *I_Ca_* augmentation, and eliminating enhanced channel recycling and resultant super-clustering. Reduced MT polymerization occurs in human heart failure where stabilized MTs form a dense network in cardiomyocytes (46, 47). The lack of polymerization and potentially enhanced MT catastrophe rates, can result in traffic jams along MTs, leading to defective cargo delivery (46). Indeed, a previous study on live ventricular myocytes reported reduced delivery of KV4.2 and KV4.3 channels to the sarcolemma due to increased MT catastrophe rates upon addition of hydrogen peroxide or in the ROS-rich post-myocardial infarction environment (48). Failing and aging myocytes display reduced Ca_V_1.2 responsivity to ISO (49, 50), thus future studies should examine whether loss of adrenergic responsivity of Ca_V_1.2 in HF occurs due to impaired channel trafficking and recycling along MTs.

Actin polymerization has also been reported to be an important determinant of cardiac ion channel trafficking, notably of Cx43 (51). Although we failed to visualize cortical F-actin because of the vast amount of sarcomeric actin in AMVMs, we found that actin disruption with lat-A led to reduced: i) ISO-stimulated mobilization of Ca_V_1.2 from EEs (Figure 3H), and ii) augmentation of Ca_V_1.2 channel expression in the PM, as indicated by superresolution GSD imaging experiments (Figure 6C) and live-cell TIRF experiments on AAV9-Ca_V_β_2a_-paGFP transduced AMVMs (Figure 4E). Furthermore, lat-A significantly blunted adrenergic regulation of the channels assessed with whole cell patch clamp (Figure 5G-I and Table 1). The degree of apparent channel insertions in response to ISO remained at a similar level to untreated cells suggesting that MTs, not actin, play the dominant role in channel delivery to the sarcolemma (Figure 4F). However, the pool of channels that was mobilized to the membrane in the presence of lat-A did not appear to belong to the EE pool, since colocalization between EEA1 and Ca_V_1.2 channels was unchanged by ISO. In the Cx43 literature, it has been proposed that Cx43 cargo on its way to the sarcolemma from the golgi along MTs, pauses at actin ‘rest-stops’ before being handed off to additional MTs to complete its journey to the membrane (51, 52). It is possible that a similar ‘rest stop’ system exists for Ca_V_1.2 channel delivery in cardiomyocytes and that lat-A mediated actin disruption and ISO-stimulation, releases this pool, allowing them to traffic to the sarcolemma along MTs. This intriguing hypothesis remains to be proven.

ISO-stimulation of AMVMs with a combined treatment with nocodazole and lat-A increased channel endocytosis as indicated by increased colocalization between EEs and Ca_V_1.2 in Airyscan images (Figure 3J), and actually led to reduced channel expression in the sarcolemma in response to ISO (Figure 4E), while adrenergic regulation of the channel was completely eliminated (Figure 5J-L). On the basis of our functional patch clamp data, it is tempting to speculate that *βAR*-mediated regulation of Ca_V_1.2 is heavily dependent on this stimulated channel recycling pathway, however, ISO-stimulation is also known to induce endocytosis and subsequent fast, actin-dependent-recycling and resensitization of the receptors themselves (53–55). It is therefore possible that the lack of functional response is simply because of a lack of sarcolemmal *βAR* expression, as internalized receptors cannot recycle back to the membrane (56). To investigate this possibility, we bypassed the receptors and stimulated adenylyl cyclase directly with forskolin and studied the effect on channel dynamics in AAV9-Ca_V_β_2a_-paGFP transduced AMVMs using TIRF imaging (Supplemental Figure 4). Forskolin (1 μM) produced a similar increase in sarcolemmal Ca_V_β_2a_-paGFP expression (21.60 ± 3.72 %) as that observed with 100 nM ISO (27.95 ± 4.31 %). Treatment of AMVMs with cytoskeletal disruptors reduced the response (Figure S4B-E). These experiments do not totally rule out a direct *βAR* effect however, as although it is GRK-mediated phosphorylation of *βARs* that increases their affinity for β-arrestin, subsequent movement into clathrin-coated pits and endocytosis; PKA-mediated desensitization of *βARs* can occur with forskolin (57). However, in a 2-3 minute exposure to forskolin or ISO, previous studies have revealed that only 20-30% of the receptors would be rendered desensitized (58), and this cannot explain our observed abolition of *βAR*-regulation of *I*_Ca_ (Figure 4), suggesting that agonist-stimulated recycling of Ca_V_1.2 is in fact a critical component of *βAR*-regulation of these channels.

After several decades of elusivity, the critical PKA phosphorylation site on the cardiac Ca_V_1.2 channel complex was recently reported to be located on Rad, a member of the Rad/Rem/Rem2/Gem/Kir (RGK) family of monomeric GTP-binding proteins that interacts with the channel via the β-subunit (5). In a disinhibition process, Rad phosphorylation is said to cause dissociation from the channel complex releasing its inhibitory hold on the channel, and unveiling the larger *I*_Ca_ we recognize as *βAR*-regulation. A Rad-mediated Ca_V_1.2 disinhibition hypothesis was also proposed several years earlier by Jonathan Satin’s group when they reported that Rad knockout mice lost adrenergic regulation (6). So how does our *βAR*-stimulated recycling of Ca_V_1.2 fit into this appealing model? We do not believe the two models are mutually exclusive but instead hypothesize that Rad inhibits Ca_V_1.2 channel function by limiting its expression at the sarcolemma, an effect that is relieved when Rad is phosphorylated by PKA. Indeed, in addition to the ability of Rad to suppress channel activity by interfering with channel *P*_o_, it has long been reported that RGK-proteins, including Rad, also reduce *I*_Ca_ by limiting Ca_V_1.2 expression at the sarcolemma (59–61). A previous study in PC12 cells reported that this effect of Rad on PM Ca_V_1.2 channel expression relied on CaM-binding to Rad, such that mutation of the CaM binding site at L281G abrogated the inhibitory effect of Rad on *I_Ca_* and increased detection of a HA-tagged Ca_V_1.2 channel at the cell surface (59). In cells transfected with WT Rad, the cell surface transport of Ca_V_1.2 was blocked. Our results in transduced AMVMs, and in tsA cells, illustrate that ISO-stimulates enhanced transport of channels to the cell surface. We speculate that this transport occurs when Rad is phosphorylated and dislodges from Ca_V_β, releasing the channel complex and allowing more of it to traffic to the surface in a Rab4a and Rab11a dependent process. Indeed, that phosphorylation of Rad may be the ‘upstream step’ that has to occur before this recycling process is initiated, explaining why overexpression of CA-Rab4a does not raise the initial expression of Ca_V_1.2 in the membrane in and of itself, but rather Rad phosphorylation must occur first. How PKA is anchored next to Rad on endosomes is another matter but may depend on the A-kinase anchoring protein D-AKAP2, which has been reported to regulate recycling of transferrin receptors via interactions with Rab4 and Rab11 (62). In line with that prediction, a human functional polymorphism in D-AKAP2 (I646V) is associated with reduced heart rate variability, indicative of a heart that cannot respond well to stressors (63). D-AKAP2 itself can be phosphorylated by PKA at residue 554 (62) and this may influence its localization. We briefly tested the hypothesis that ISO-stimulation of *βARs* would promote enhanced colocalization between D-AKAP2 and Rab11-positive endosomes finding an enhanced association (Figure S5). While this is admittedly, a correlative result, resolving these mechanistic details will make for an interesting future project.

Overall, our data indicate that there is an endosomal pool of intracellular Ca_V_1.2 channels that is stimulated to recycle to the sarcolemma upon *βAR*-activation. Both Rab4a-mediated fast recycling pathways, and Rab11a-slow recycling pathways contribute to the response. Recycling channels insert into the sarcolemma by traveling along MTs likely anchored at the PM by BIN1. Cytoskeletal disruption creates traffic jams in the endosomal pathway and impedes the recycling response. Finally, electrophysiology data indicates that *βAR*-regulation of cardiac Ca_V_1.2 channel function is abolished by treatments that interfere with this stimulated recycling pathway. Collectively, these results suggest that cardiomyocytes have an endosomal reservoir of Ca_V_1.2 channels that is rapidly mobilized to the sarcolemma in response to *βAR*-stimulation, and that this stimulated insertion is fundamentally required for *βAR*-regulation of these channels.

## Materials and Methods

Detailed methods can be found in the SI Appendix. Briefly, AMVMs were enzymatically isolated using standard Langendorff technique as described previously (23, 64, 65). Fixed, immunostained AMVMs were imaged on a Zeiss Airyscan confocal microscope (as described in (66, 67)) or a Leica 3D-GSD-SR microscope to assess the distribution of Ca_V_1.2 channels and various endosome populations (Airyscan only for the latter). Live cell TIRF imaging experiments of AAV9-Ca_V_β_2a_-paGFP transduced AMVMs or transiently transfected tsA-201 cells were performed on an Olympus IX-83 inverted microscope with a Cell-TIRF MITICO module. Live cell 4D-imaging of transduced AMVMs was performed at 37 °C on an Andor W-1 Spinning Disk confocal microscope. Rab mutant plasmids used for transient transfection of tsA-201 cells were gifts from Dr. Nipavan Chiamvimonvat (UC Davis, Davis, CA, USA) and Dr. Jose A. Esteban (*Centro de Biología Molecular ‘Severo Ochoa’* (*CSIC-UAM*), Madrid, Spain).

## Acknowledgments

We wish to thank Dr. Luis Fernando Santana for the use of his GSD microscope, and for reading and commenting on this manuscript along with Dr. Manuel F. Navedo, and Dr. Maartje Westhoff.

