## Supplemental Information Appendix for "β-adrenergic control of sarcolemmal Ca_V_1.2 abundance by small GTPase Rab proteins"

**SI Appendix**

### Detailed Materials and Methods

#### Ethical Approval

All animal procedures described in this study were reviewed and approved by the University of California Davis Institutional Animal Care and Use Committee (IACUC) and conducted in accordance with the Guide for the Care and Use of Laboratory Animals (1).

#### Isolation of mouse ventricular myocytes

Ventricular myocytes were isolated from male and female C57BL/6J mice purchased from The Jackson Laboratory (Sacramento, CA, USA). 12-20 week old mice were euthanized by intraperitoneal injection of pentobarbital solution (> 100 mg/kg; Beuthanasia-D Special; Merck Animal Health, Madison, NJ, USA). The heart was removed via sharp dissection and placed in a 150 µM EGTA supplemented ice-cold digestion buffer containing: 130 mM NaCl, 5 mM KCl, 3 mM Na-pyruvate, 25 mM HEPES, 0.5 mM $MgCl_2$, 0.33 mM $NaH_2PO_4$ and 22 mM glucose. Myocytes were isolated via enzymatic digestion of the heart using the Langendorff technique as described previously (2-4). Briefly, the aorta was cannulated and the heart perfused with warmed 37°C digestion buffer until it ran clear of blood. The perfusing solution was then switched to digestion buffer containing 50 µM $CaCl_2$ (ThermoFisher), 0.04 mg/ml protease and 1.4 mg/ml type 2 collagenase (Worthington Biochemical, Lakewood, NJ, USA) until digestion was complete (as assessed by appearance and firmness of the heart). The atria were removed, and ventricles were sliced into smaller pieces, placed in warmed digestion buffer supplemented with 0.96 mg/mL collagenase, 0.04 mg/mL protease, 100 µM $CaCl_2$ and 10 mg/ml BSA, and a transfer pipette was used to gently separate the tissue into individual cells. Cardiomyocytes were separated from the digestion buffer by gravity pelleting for 20 min at room temperature. The supernatant was discarded and the pellet resuspended in a washing buffer consisting of digestion buffer supplemented with 10 mg/ml BSA and 250 µM $CaCl_2$. Cells were gravity pelleted once more for 20 mins at room temperature, the wash buffer supernatant was discarded, and the cells were finally suspended in the appropriate solution for the desired experiments.

#### Immunostaining and Airyscan microscopy

Fresh ventricular myocytes were plated onto poly-L-lysine (0.01%; Sigma-Aldrich) and laminin (20 µg $mL^{-1}$; Life Technologies, Carlsbad, CA, USA) coated glass coverslips and allowed to adhere for 45 mins at room temperature. Myocytes were washed with PBS and before a 3 min treatment with either PBS (control) or 100 nM isoproterenol (ISO). In experiments focused on the effects of cytoskeletal disruption, myocytes first underwent a 2 h room temperature incubation in either: a) 10 µM nocodazole; b) 5 µM latrunculin-A (lat-A) or c) both drugs in combination (combo); before proceeding to 3 min control or ISO stimulation phase. Myocytes were then fixed in a 4% paraformaldehyde solution with a cytoskeletal preserving buffer (PEM) containing 80 mM PIPES, 5 mM EGTA, and 2 mM $MgCl_2$ at 37°C for 10 min. Cells were rinsed with PBS and incubated for 10 mins in 0.5% Triton-X 100 (Sigma-Aldrich) prior to an overnight step at 4°C in blocking solution containing 50% SEA Block (Thermo Fisher Scientific, Rockford, IL, USA) and 0.5% v/v Triton X-100 in PBS.

Primary antibodies were diluted in blocking solution and applied to coverslip adherent cells for 45 min at room temperature. For double and triple labeling, two or three primary antibodies were used simultaneously. The following primary antibodies were used: rabbit polyclonal IgG anti-$Ca_v$1.2 (1:300; Alomone labs ACC-003), mouse monoclonal anti-EEA1 IgG1 (1:250; Abcam ab70521), mouse monoclonal $IgG_{2a}$ anti-Rab4 (1:250; Santa Cruz sc-376243), mouse monoclonal $IgG_1$ anti-Rab11a (1:250; Santa Cruz sc-166523), mouse monoclonal $IgG_1$ anti-Rab7 (1:250; Santa Cruz sc-376362), rat $IgG_{2a}$ anti-α tubulin (1:100 Thermofisher MA1-80017), and mouse monoclonal $IgG_{2a}$ anti-AKAP10 (1:100; Abnova H00011216-M04). After thorough washing with PBS, cells were incubated in Alexa Fluor conjugated secondary antibodies (1:1000; Life Technologies) or Phalloidin (0.6 U) for 30 min at RT. The following secondary antibodies were used: Alexa Fluor 488 goat anti-rabbit, Alexa Fluor 568 goat anti-mouse $IgG_1$, Alexa Fluor 568 goat anti-Rabbit IgG, Alexa Fluor 647 goat anti-mouse $IgG_{2a}$, goat anti-rat $IgG_{2a}$-FITC, and Phalloidin 647 nm (Life Technologies). After thorough washing in PBS, coverslips were mounted onto glass slides and examined on a Zeiss LSM 880 super-resolution microscope equipped with an Airyscan detector and a Plan-Apochromat 63×/1.40 oil DIC M27 objective. Images were acquired using Zen software.

**tsA-201 cell culture and transfection**

tsA-201 cells (Sigma–Aldrich, St. Louis, MO) were maintained in Dulbecco's Modified Eagle's medium (DMEM) supplemented with 10% fetal bovine serum (FBS) and 1% penicillin/streptomycin in a cell culture incubator (37°C, 95:5% $O_2$:$CO_2$). Cells were passaged every 3-4 days and transfected at ~70% confluence using JetPRIME DNA Transfection reagent

(Polyplus Transfection, New York, NY, USA). Cells were plated onto glass coverslips at low density after 6 h of transfection and used for experiments the following day ~16 h later.

**Plasmid and viral constructs**

For expression of $Ca_V1.2$ in tsA201 cells, 1000 ng of each of the following plasmid constructs were transfected: rabbit cardiac isoform of $Ca_v1.2$ $\alpha_{1c}$ (GenBank accession number: NP_001129994.1; provided by William Catterall, University of Washington, Seattle, WA, USA), rat $Ca_V\alpha_2\delta$ (AF286488), rat $Ca_V\beta_3$ (M88751; both auxiliary subunits were provided by Dr. Diane Lipscombe; Brown University, Providence, RI, USA). The pore-forming $\alpha_{1c}$ construct was tagged at its C-terminus with a monomeric GFP(A206K) or a tagRFP to permit confirmation of transfection and to monitor channel localization. In experiments designed to interrogate the role of Rab4a and Rab11a in ISO-stimulated $Ca_V1.2$ channel recycling, cells were co-transfected with pEGFP-Rab plasmids with previously described single amino acid substitutions conferring either constitutive activity (CA; GTP bound/locked) or lack of function (dominant negative, DN; GDP-bound/locked). Accordingly, CA-Rab4a(Q67L), DN-Rab4a(S22N), DN-Rab11a(S25N) were provided by Dr. Nipavan Chiamvimonvat (UC Davis, Davis, CA, USA) and Dr. Jose A. Esteban (*Centro de Biología Molecular 'Severo Ochoa' (CSIC-UAM)*, Madrid, Spain).

**TIRF imaging of transduced mouse ventricular myocytes and tsA-201 cells**

Three-week old mice under isoflurane anesthesia were retro-orbitally injected with AAV9-$Ca_V\beta_{2a}$-paGFP. 7-12 weeks later, the mice were euthanized and individual ventricular myocytes were isolated from the transduced heart using the method described above. Isolated cells were allowed to adhere to poly-L-lysine coated glass coverslips for 15 mins prior to imaging. In experiments focused on the effects of cytoskeletal disruption, myocytes first underwent a 2 h room temperature incubation in either: a) 10 µM nocodazole; b) 5 µM lat-A, or c) combo; before imaging. In TIRF experiments, an Olympus IX83 inverted microscope equipped with a Cell-TIRF MITICO and a 60×/1.49 NA TIRF objective lens was used to image the attached myocytes. 405 nm epi-fluorescent light was used to photoactivate the paGFP on the transduced paGFP-$Ca_V\beta_{2a}$, 488 nm laser light was then used to excite activated GFP. Myocytes were constantly perfused with Tyrode's solution (140 mM NaCl, 5 mM KCl, 10 mM HEPES, 10 mM glucose, 2 mM $CaCl_2$ and 1 mM $MgCl_2$; pH adjusted to 7.4 with NaOH) and images were acquired at 10.34 FPS. After a 300 frame control period, 100 nM ISO, or 1 µM forskolin was perfused for 3 min (1861 frames).

In tsA-201 cell experiments, cells perfused with Ringer's solution (160 mM NaCl, 2.5 mM KCl, 2 mM $CaCl_2$, 1mM $MgCl_2$, 10 mM HEPES and 8 mM glucose; pH adjusted to 7.4 with NaOH). were imaged on the same system using a 100x/1.49 NA TIRF objective.

ImageJ/FIJI was used for image analysis. Images stacks were subjected to bleach correction, a 20-pixel rolling ball background subtraction and a 10-frame moving average. To observe and quantify the recycled, endocytosed and steady population of $Ca_v\beta_{2a}$-paGFP, maximum intensity z-projections of the 300 frame control period and the final 300 frames of the ISO period were generated, thresholded and made binary. Next, using the 'image calculator' function in FIJI, to generate a map of channels that were: i) inserted/recycled into the membrane during ISO, the binary control image was subtracted from the binary ISO image, ii) endocytosed during ISO, the binary ISO image was subtracted from the binary control image, and iii) static, population image the binary control and ISO images were multiplied. A custom written code was written in Matlab to automatically identify ROIs where channels were inserted or removed during the experiment, and to plot the time course of fluorescence intensity in those regions. We have previously described the use of this code for the analysis of $Ca^{2+}$ sparklets and it is freely available with that previous publication (3).

For experiments conducted at 37°C, coverslip adherent myocytes were imaged on an Olympus IX83 inverted microscope equipped with an Andor (Belfast, Northern Ireland) W1-spinning disk and a 60×/1.49 NA TIRF objective lens. Physiological temperature was maintained using a stage-top incubator (Okolab USA Inc., Ambridge, PA).

**Whole cell patch clamp electrophysiology**

Whole cell $Ca_v$1.2 channel currents ($I_{Ca}$) were recorded from freshly isolated ventricular myocytes using fire polished borosilicate glass pipettes with 1-3 MΩ resistance, filled with an internal solution containing 87 mM Cs-aspartate, 20 mM CsCl, 1 mM $MgCl_2$, 10 mM HEPES, 10 mM EGTA and 5 mM MgATP (pH adjusted to 7.2 with CsOH). Myocytes were initially perfused with an external Tyrode's solution containing 140 mM NaCl, 5 mM KCl, 10 mM HEPES, 10 mM glucose, 2 mM $CaCl_2$ and 1 mM $MgCl_2$ (pH adjusted to 7.4 with NaOH). Upon attainment of whole cell configuration, the external solution was switched to one containing 5 mM CsCl,10 mM HEPES,10 mM Glucose, 140 mM NMDG, 1 mM $MgCl_2$ and 2 mM $CaCl_2$ (pH adjusted to 7.3 with HCl). Cells remained at a holding potential of -80 mV for 8 mins prior to the recording of the control currents to minimize the effects of current run-down on our control vs ISO comparisons. Currents were subsequently elicited using an established protocol, briefly, myocytes were held at -80 mV and stepped initially to -40 mV for 100 ms (to inactivate $Na^+$ channels), followed by 300 ms steps to test potentials ranging from -60 - +80 mV. Currents were sampled at a frequency of 10 kHz, and low-pass–filtered at 2 kHz using an Axopatch 200B amplifier (Molecular Devices, Sunnyvale, CA, USA), digitized using a Digidata 1550B plus Humsilencer (Molecular Devices) and acquired using pClamp, version 10.7 (Molecular Devices). Analysis was performed with Clampfit software (Molecular Devices) and current-voltage relationship were plotted using Prism (GraphPad Software Inc., La Jolla, CA, USA). Membrane potentials were corrected for liquid junction potential.

In experiments designed to evaluate the functional effects of cytoskeletal disruption, isolated myocytes were patched after 2 h room temperature incubations in cytoskeletal disruptors as outlined above.

**Ground State Depletion Super-Resolution Microscopy**

Coverslips (No. 1.5) were sonicated for 20 mins in 2 N NaOH prior to use to remove any contaminants and washed thoroughly from base with several rinses in de-ionized water. Isolated AMVMs were plated onto poly-L-lysine and laminin-coated coverslips and placed in a 37°C incubator to adhere for 45 min. Adherent AMVMs were next washed with PBS ahead of an 8 min treatment with either PBS (control) or 100 nM ISO. As with the Airyscan experiments described above, some myocytes first underwent a 2 h room temperature incubation in either: a) 10 µM nocodazole; b) 5 µM lat-A or c) combo. AMVMs were then fixed in ice-cold 100% methanol (Fisher Scientific) for 5 min at -20°C, before thorough washing in PBS, and blocking for 1 h at room temperature in 50% SEA Block (ThermoFisher) and 0.5% v/v Triton X-100 (Sigma-Aldrich) in PBS. Primary antibody incubation was performed overnight at 4°C in rabbit polyclonal IgG anti-$Ca_V$1.2 (Alomone labs ACC-003) diluted 1:300 in blocking buffer (20% SEA BLOCK, 0.5% Triton X-100). The next day, excess primary antibody was washed off with PBS, and secondary antibody incubation commenced for 1 h at room temperature with Alexa Fluor 647-conjugated donkey anti-rabbit (Life Technologies) diluted 1:1000 in PBS. After a final series of washes in PBS, the coverslips were then mounted onto glass depression slides (neoLab, Heidelberg, Germany) with a cysteamine (MEA)-catalase/glucose/glucose oxidase (GLOX) imaging buffer containing TN buffer (50 mM Tris pH 8.0, 10 mM NaCl), a GLOX oxygen scavenging system (0.56 mg $mL^-$

[1] glucose oxidase, 34 µg $mL^{-1}$ catalase, 10% w/v glucose) and 100 mM MEA. Twinsil dental glue (Picodent, Wipperfürth, Germany) and aluminum tape (T205-1.0 - AT205; Thorlabs Inc., Newton, NJ, USA) were used to hold the coverslip in place on the slide and to exclude oxygen. AMVMs were imaged on a Leica 3D-GSD-SR microscope (Leica Microsystems, Wetzlar, Germany) in TIRF mode with a penetration depth of 150 nm, using an oil-immersion HC PL APO 160 ×/1.43 NA TIRF objective (Leica) as previously described (5). Ground state depletion was performed and dye-blinking was elicited using a 642 nm/500 mW laser, back-pumping was performed with a high-energy 405 nm/30 mW laser. Photon emission was detected with an iXon3 EM-CCD camera (Andor). Raw blinking images were collected at 100 fps for 38,000 frames using Leica Application Suite software and visualized in a live localization map. $Ca_V1.2$ cluster areas and total integrated density of localization maps with a 10 nm pixel, were determined in ImageJ/Fiji.

**Chemicals and statistical analysis**

Chemical reagents were obtained from Sigma-Aldrich unless otherwise stated. Data are reported as the mean ± SEM. *N* reflects the number of animals used in a dataset, whereas *n* reflects the number of cells. Student's *t* test was used for paired datasets comparison if the data passed a normality test. If not, a non-parametric test was performed. Two-way ANOVAs were used to compare data sets with two independent variables, eg. to test whether sex and ISO affects $Ca_V1.2$ and EEA1 colocalization. $P < 0.05$ was considered statistically significant ($^{*}$ $P < 0.05$; $^{**}$ $P < 0.01$; $^{***}$ $P < 0.001$).

### Supplemental Figures

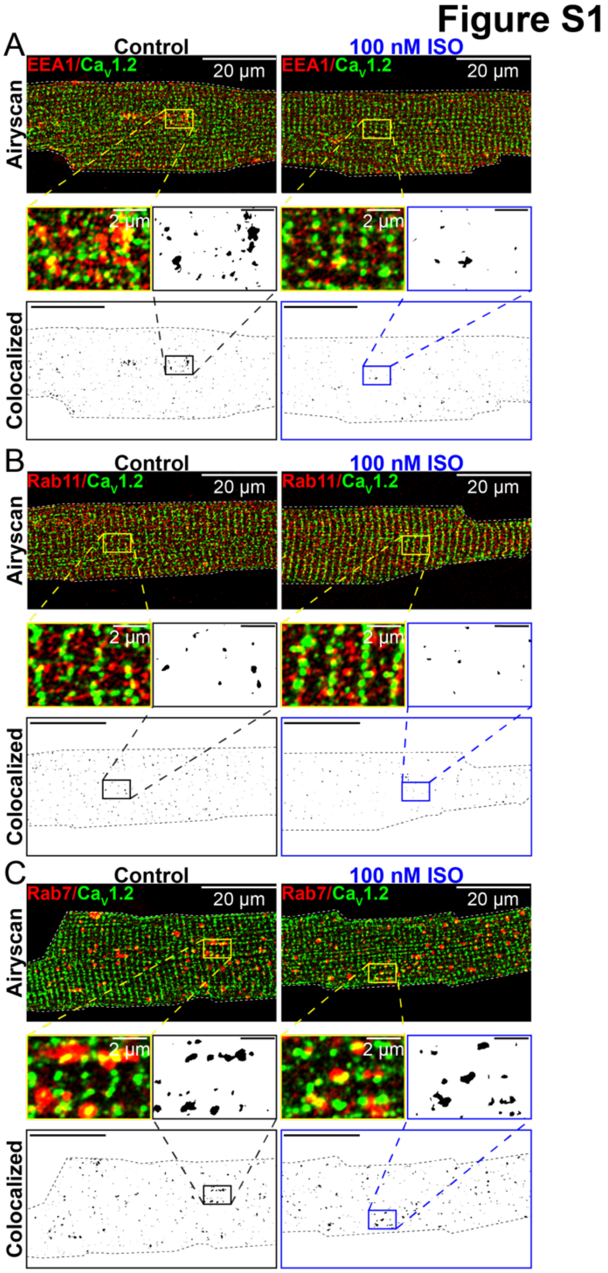


**Figure S1. A pool of intracellular $Ca_V1.2$ channels resides on endosomes in female AMVMs.** Two-color Airyscan super-resolution images of control and 100 nM ISO-stimulated, female AMVMs immunostained to examine $Ca_V1.2$ localization on: (*A*) EEA1 positive EEs, (*B*) Rab11 positive REs, and (*C*) Rab 7 positive LEs and lysosomes. In each case, binary colocalization maps (*bottom*) display pixels in which $Ca_V1.2$ and endosomal expression precisely overlapped.

### Figure S2

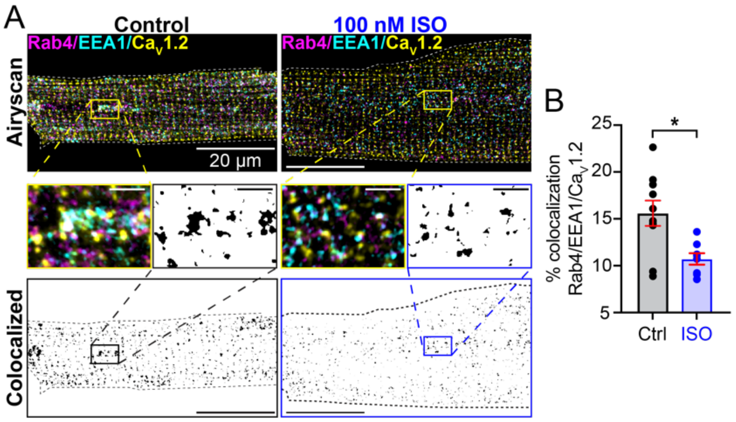


**Figure S2. A Ca$_V$1.2 channel pool on EEA1/Rab4 positive endosomes.**

(*A*) Three-color Airyscan super-resolution images of control (*left*; $N = 2$, $n = 10$), and 100 nM ISO-stimulated (*right*; $N = 2$, $n = 8$) male AMVMs immunostained to examine distributions of Ca$_V$1.2 and early endosomes labeled with anti-EEA1 and anti-Rab4. *Bottom*: binary colocalization maps display pixels in which Ca$_V$1.2, EEA1, and Rab4 all precisely overlapped. (*B*) Histogram summarizing the % colocalization between all three labels, Ca$_V$1.2, EEA1, and Rab4.

### Figure S3

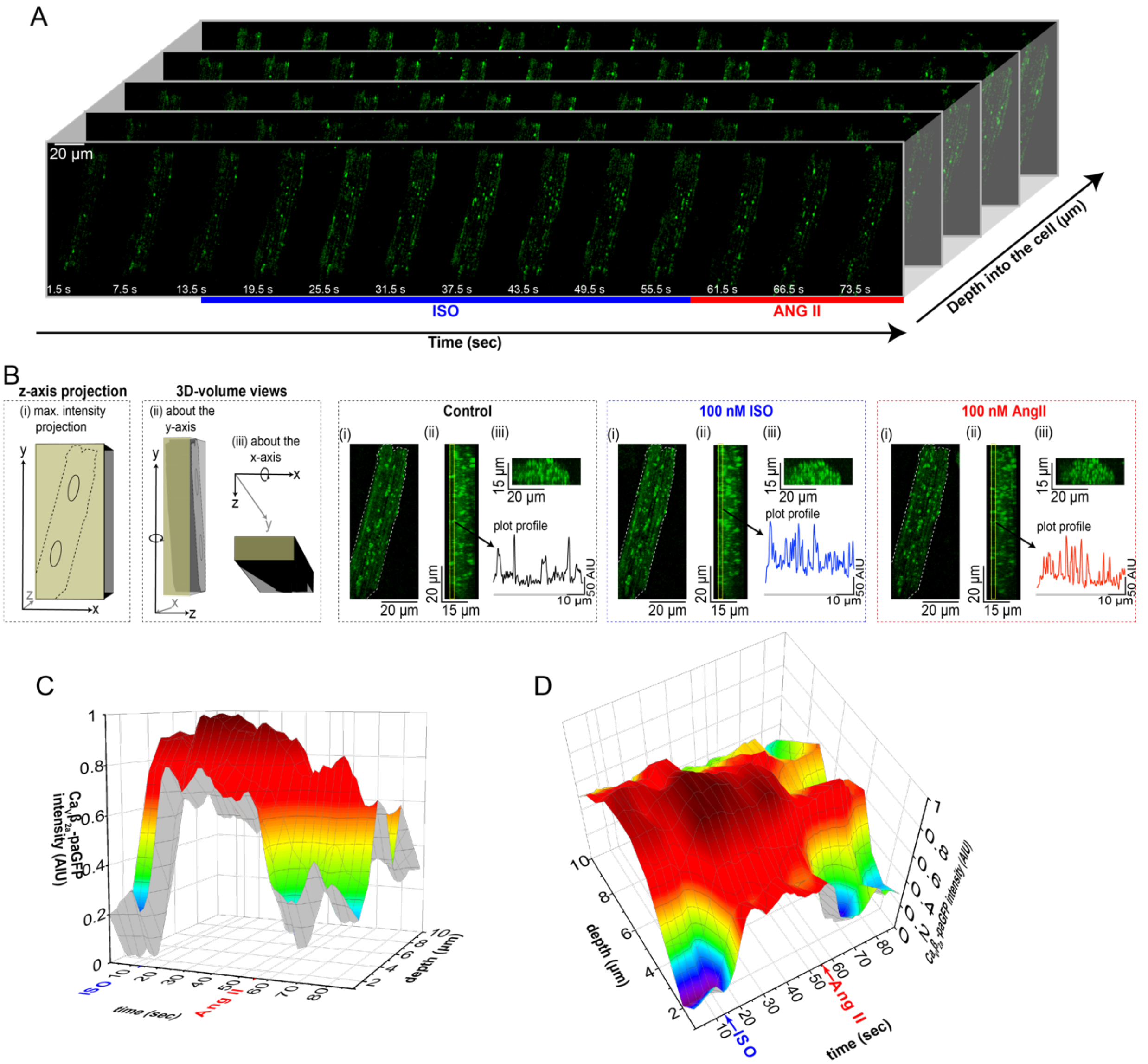


**Figure S3. Rapid ISO-stimulated channel recycling at physiological temperature.**

4D-spinning disk confocal images of AAV9-Ca$_v\beta_{2a}$-paGFP transduced AMVMs maintained at 37°C before and after 100 nM ISO and 100 nM Angiotensin II (AngII) treatment. (*A*) is an illustration of the experimental design. Briefly, a z-stack was performed through the depth of the cell. This was then repeated over the course of a time series during which 100 nM ISO was perfused on, followed by 100 nM AngII. (*B*) Schematic describing the 3 projections seen in the next three panels: (*i*) a max intensity projection through the depth (z-axis) of the cell, (*ii*) a 3D-volume view about the y axis and (*iii*) a 3D-volume view about the x-axis of the cell. Images are shown of each of these projections during control, ISO and AngII perfusion periods. Vertical plot profiles are shown in each case for the yellow box ROI outlined in (*ii*) and shows the intensity of $Ca_v\beta_{2a}$-paGFP at the cell surface. ISO was observed to trigger the movement of more $Ca_v\beta_{2a}$-paGFP from within the cell to the surface, while AngII treatment stimulated the channels to move back into positions, deeper within the cell. (*C*) and (*D*) show 3D plots of the results from the cell shown in A and B. Warmer colors indicate higher intensity of $Ca_v\beta_{2a}$-paGFP. Note the red color showing high intensity $Ca_v\beta_{2a}$-paGFP emission propagates quickly ($\tau$ = 2.53 $\pm$ 0.28 sec) toward the cell surface in response to ISO, reaching a plateau until 100 nM ISO was added, causing the surface intensity of $Ca_v\beta_{2a}$-paGFP emission to drop almost as rapidly ($\tau$ = 2.94 $\pm$ 0.96 sec). Note the units here are seconds, whereas those at room temperature were minutes.

**Figure S4**

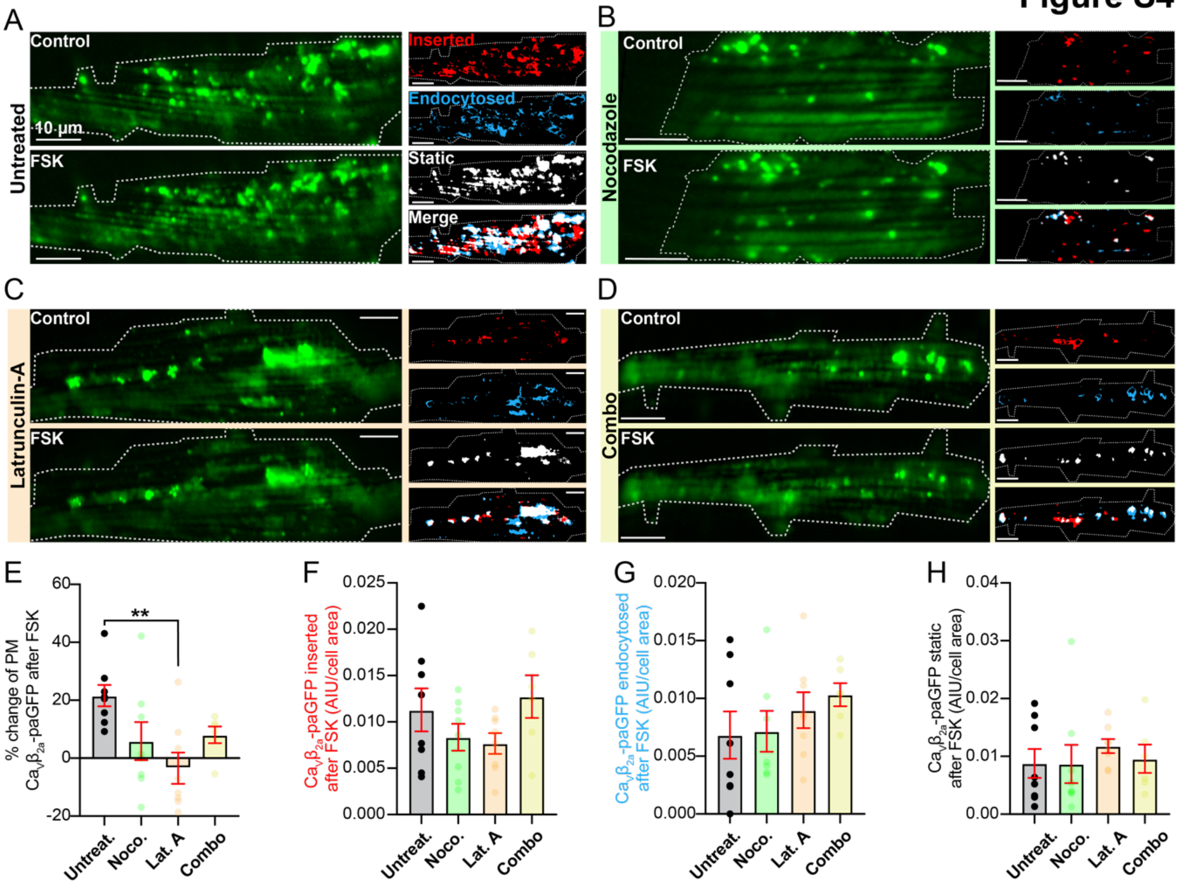


**Figure S4. Stimulated $Ca_v\beta_{2a}$-paGFP recycling also occurs with forskolin.**

(*A*) TIRF images of GFP fluorescence emission from $Ca_v\beta_{2a}$-paGFP transduced cardiomyocytes (*N* = 3, *n* = 8) before (*top*) and after 1 µM FSK (*bottom*). Same format for cells pre-treated with (*B*) nocodazole (*N* = 2, *n* = 8), (*C*) lat-A (*N* = 2, *n* = 8), and (*D*) a combination of nocodazole and lat-A (*N* = 2, *n* = 6). (*E-H*) Histograms summarizing statistics for these experiments.

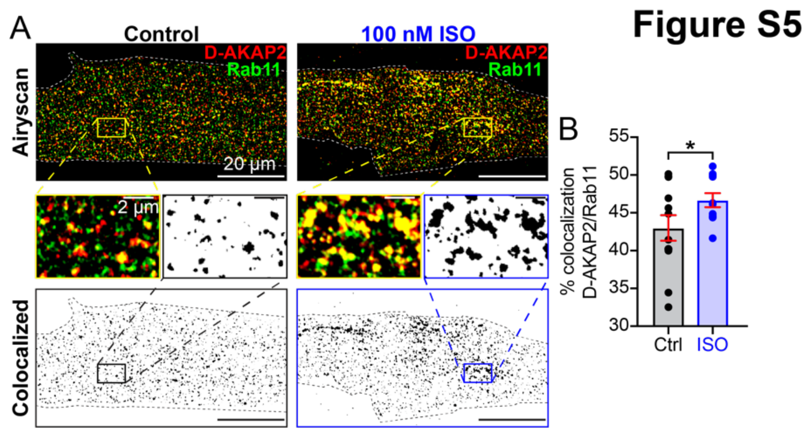


**Figure S5. ISO-stimulated recruitment of D-AKAP2 to Rab11-positive REs.** (*A*) Two-color Airyscan super-resolution images of control (*left; N* = 2, *n* = 12) and 100 nM ISO-stimulated (*right; N* = 2, *n* = 11), AMVMs immunostained to examine D-AKAP2 localization on Rab11-positive REs. Binary colocalization maps (*bottom*) display pixels in which D-AKAP2 and Rab11 expression precisely overlapped. (*B*) Histogram summarizing the % colocalization between D-AKAP2 and Rab11 ($P$ = 0.038).

**Movie Legends**

**Movie 1. ISO-stimulated channel recycling.** TIRF time series obtained from an untreated AAV9-$Ca_V\beta_{2a}$-paGFP transduced ventricular myocyte. *β*AR activation with 100 nM ISO stimulated dynamic augmentation of $Ca_V$1.2 channel insertions in the sarcolemma as indicated by the $Ca_V\beta_{2a}$-paGFP channel proxy.

**Movie 2. Microtubules provide delivery pathways for recycling $Ca_V$1.2 channels.** TIRF time series obtained from a nocodazole-treated (10 µM for 2 h), AAV9-$Ca_V\beta_{2a}$-paGFP transduced ventricular myocyte. *β*AR activation with 100 nM ISO stimulated conspicuously fewer sarcolemmal channel insertions than that observed in control untreated cells.

**Movie 3. F-actin is an important cytoskeletal component in ISO-stimulated augmentation of sarcolemmal $Ca_V$1.2.** TIRF time series obtained from a latrunculin-A-treated (5 µM for 2 h), AAV9-$Ca_V\beta_{2a}$-paGFP transduced ventricular myocyte. *β*AR activation with 100 nM ISO failed to trigger stimulated augmentation of sarcolemmal $Ca_V$1.2.

**Movie 4. F-actin and microtubules are important conduits of ISO-stimulated $Ca_V$1.2 trafficking.** TIRF time series obtained from an AAV9-$Ca_V\beta_{2a}$-paGFP transduced ventricular myocyte, treated for 2 h with nocodazole (10 µM) and latrunculin-A (5 µM). *β*AR activation with 100 nM ISO failed to trigger stimulated augmentation of sarcolemmal $Ca_V$1.2 indicating the importance of these cytoskeletal elements in this dynamic trafficking response.
